# Disassembly of ALS condensates by a disordered peptide

**DOI:** 10.1101/2025.03.19.643735

**Authors:** Mani Garg, Nayana Vinayan, Kumkum Nag, Riya Dhage, Poulami Ghosh, Kesavardhana Sannula, Purusharth I Rajyaguru

**Affiliations:** Department of Biochemistry, Indian Institute of Science, Bangalore 560012

**Keywords:** Neurodegenerative disorder, amyotrophic lateral sclerosis (ALS), ribonucleoprotein (RNP) condensates, condensate disassembly, intrinsically disordered region (IDR), low-complexity sequences (LCSs), RGG motif, Sbp1, FUS, FUS-P525L, TDP43, TDP43-ΔNLS

## Abstract

Mislocalization and cytoplasmic condensation of nuclear proteins, such as FUS and TDP43, are key features of amyotrophic lateral sclerosis (ALS) and frontotemporal dementia (FTD). However, mechanisms governing the disassembly of these pathological condensates remain elusive. Here, we identify a disordered RGG peptide as a disassembly factor for mutant FUS and TDP43 condensates. Using yeast, mammalian cells, and *in vitro* assays, we observe that an RGG motif reduces condensate burden and toxicity while restoring nuclear localization and RNA regulatory defects associated with the FUS-P525L mutant. The RGG motif interacts with FUS, competitively inhibiting FUS self-association. Overall, our findings uncover a direct role for a disordered RGG motif in the disassembly of ALS-associated condensates, highlighting the possibility of a new therapeutic for post-symptomatic ALS.

## Introduction

The assembly and disassembly of RNP condensates (such as P-bodies and stress granules) are tightly regulated to maintain cellular homeostasis. While the formation of stress granules (SGs) is essential for many cellular responses^1–3^, a timely disassembly is also important for maintaining cellular health. Persistent or aberrant SGs are increasingly implicated in neurodegenerative disorders, including amyotrophic lateral sclerosis (ALS) and frontotemporal dementia (FTD)^4^. In such cases, SGs often undergo a liquid-to-solid transition, promoting the entrapment of nuclear RNA-binding proteins such as FUS and TDP43 within cytoplasmic condensates. These mislocalized proteins contribute to disease by both gaining toxic function and losing essential nuclear roles. Although previous studies have identified factors that either limit the recruitment of FUS and TDP43 to SGs or modulate their toxicity^5–9^, the disassembly of pre-existing pathogenic condensates remains an underexplored therapeutic avenue. Given that most ALS cases are sporadic and diagnosed after symptoms manifest, an effective therapeutic strategy would be to focus on slowing disease progression by targeting disassembly. In this direction, the identification of disassembly factors will be beneficial for these conditions where the proteins are already mis-localized to these condensates.

Intrinsically disordered regions (IDRs) containing proteins with additional folded RNA-binding domains have been extensively associated with RNP condensate assembly^1,10^. IDRs are often characterized by low-complexity sequences (LCS) consisting of repeats of charged amino acids such as arginine and glycine. These sequences can participate in multiple low-affinity interactions, such as pi-pi and cation-pi interactions^11^. Often, the repeats are interspersed with aromatic amino acids such as phenylalanine and tyrosine, which further facilitate cation-pi as well as pi-pi interactions. Moreover, the disorder provides accessibility of various residues to post-translational modifications (PTMs), such as tyrosine phosphorylation and arginine methylation, and readily allows interactions with other ligands^12^. We hypothesised that proteins with LCS, such as those with RGG motifs, could participate in multiple low-affinity interactions that affect the pre-formed pathological assemblies of FUS and TDP43 implicated in ALS. We report here the role of the RGG motif from Sbp1, a yeast protein, to test its impact on FUS and TDP43 condensates. Sbp1 harbors a central disordered arginine and glycine (RGG/RG) repeat rich RGG-motif flanked on either side by RNA-recognition motifs (RRM, Figure 1A and Supplementary Figure 1A). Our data from *Saccharomyces cerevisiae*, mammalian cells, and *in vitro* experiments demonstrate that RGG peptides can disassemble pathological condensates of both FUS and TDP43, raising the possibility of a therapeutic potential for RGG peptides in treating ALS.

**Figure 1:**
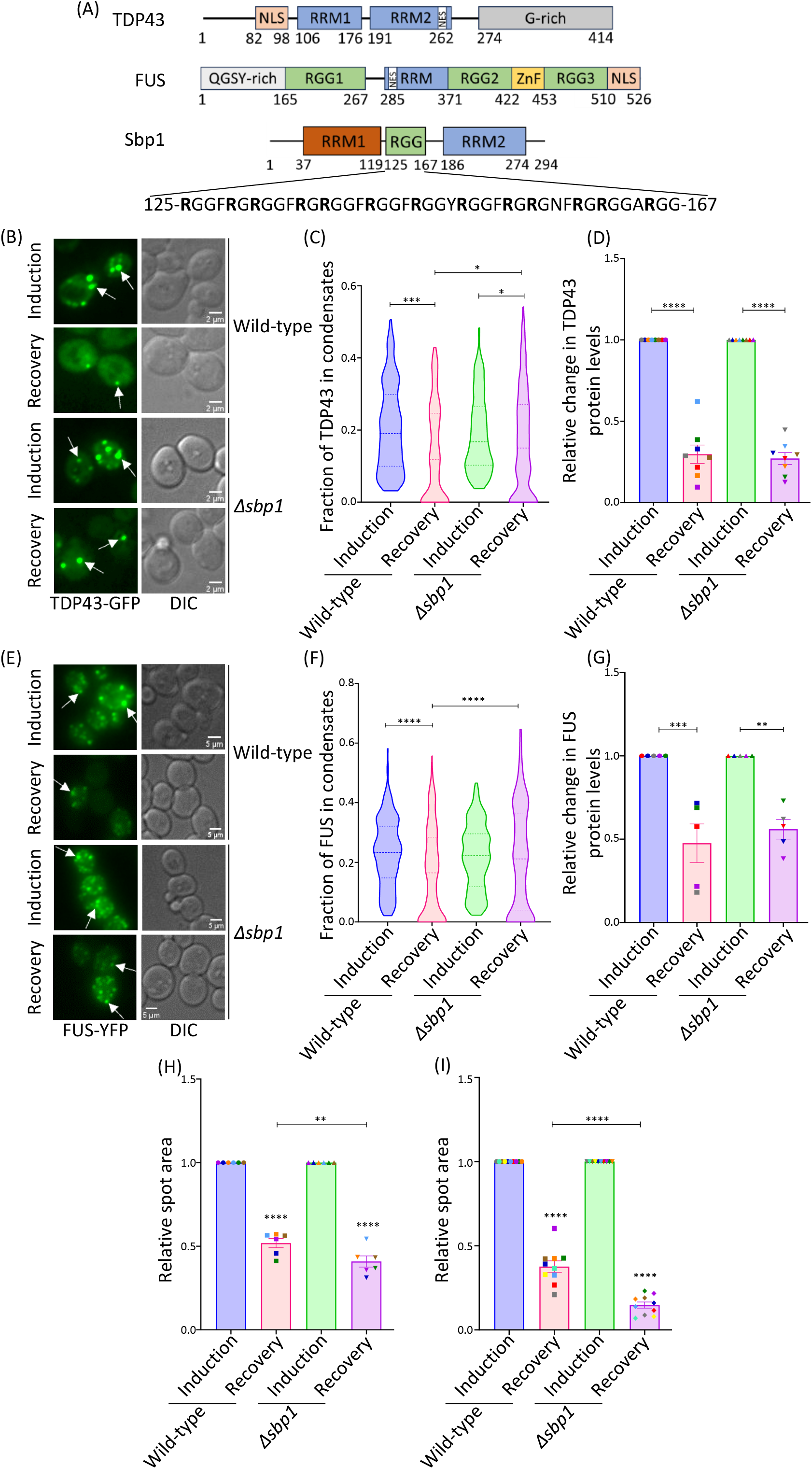
Δsbp1 cells are defective in the disassembly of TDP43 and FUS condensates in yeast and show enhanced toxicity. (A) Schematic representation of TDP43, FUS, and Sbp1 proteins. For Sbp1, the sequence architecture of the RGG-motif is also presented. (B) Representative images for the induction and recovery phases of the wild-type and Δ*sbp1* cells transformed with the Gal-*TDP43-GFP* plasmid. White arrows mark the presence of TDP43 condensates. Scale bar=2um. (C) Graph representing the fraction of the TDP43 protein present in condensates per cell. A minimum of 50 cells per experiment were analyzed from 4 independent experiments (n=4) as performed in B. An unpaired t-test was used to calculate the significance. (D) Graph depicting the relative change in TDP43 protein levels during the recovery phase as compared to the respective induction condition. Significance was calculated using Tukey’s multiple comparisons test (2-way ANOVA, n=8). (E) Representative images for the induction and recovery phases of wild-type and Δ*sbp1* cells transformed with the Gal-*FUS-YFP* plasmid. White arrows mark the presence of FUS condensates. Scale bar=5um. (F) Graph representing the fraction of the FUS protein in condensates per cell. A minimum of 50 cells per experiment were analyzed from 5 independent experiments as performed in E (n=5). An unpaired t-test was used to calculate the significance. (G) Graph depicting the relative change in FUS protein levels during the recovery phase compared to the respective induction condition. Significance was calculated using a Tukey’s multiple comparisons test (2-way ANOVA, n=5). (H and I) Graphs representing the yeast cell viability as calculated from the spotting assays of wild-type and Δ*sbp1* cells transformed with either Gal-*TDP43-GFP* (H) or Gal-*FUS-YFP* (I) expressing constructs. Values were normalized with respect to their empty vector controls. Significance was calculated by Tukey’s multiple comparisons test (2-way ANOVA; n=6 and n=10 for H and I, respectively). Error bars in all graphs represent mean ±SEM, and the same color points depict the data from a single experimental set for all the graphs. *, **, ***, and **** denote p-value≤0.05, ≤0.01, ≤0.001, and ≤0.0001, respectively.

## Results

### TDP43 and FUS condensates accumulate in Δ*sbp1*

*Saccharomyces cerevisiae* has been extensively used as a model to understand various neurological disorders^13–19^. Overexpression of ALS-associated proteins, such as TDP43 and FUS, using a galactose-inducible promoter, is known to be cytotoxic in yeast, and several genetic modifiers of this toxicity have been identified^13,20–27^. Leveraging this model, we sought to examine the role of the RGG motif-containing protein Sbp1 in the disassembly and toxicity of TDP43 and FUS condensates. Yeast cells overexpressing TDP43 and FUS under a galactose-inducible promoter were incubated in galactose-containing medium to induce the expression of each protein. During this phase, the newly translated TDP43 and FUS proteins induce stress (termed ‘induction’) and accumulate in the cytoplasmic condensates (Figure 1B and E). To initiate disassembly, the cells were then allowed to grow in glucose-containing medium so that TDP43 and FUS levels were reduced because of the inhibition of the galactose promoter (Figure 1D and G). This reduction leads to the rescue of cells from stress (termed ‘recovery’) and a subsequent decrease in the number of condensates. The amount of protein present in the condensates was then assessed and compared between wild-type and Δ*sbp1* cells to dissect the role of Sbp1 protein in the assembly and disassembly of the TDP43/FUS condensates.

The induction of TDP43 condensates was first assessed in wild-type and Δ*sbp1* cells. The fraction of protein localized to the condensates was comparable in both genotypes after induction (Figure 1B and C). However, during the recovery phase, Δ*sbp1* cells exhibited a markedly higher fraction of TDP43 retained in condensates compared to wild-type cells, suggesting a defect in condensate disassembly (Figure 1B and C). The dynamics of FUS condensates also followed a similar trend in the Δ*sbp1* as compared to the wild-type background (Figure 1E and F). While the fraction of FUS protein localized to condensates upon stress was comparable between wild type and Δ*sbp1*, there was significantly more FUS protein in condensates of Δ*sbp1* as compared to the wild-type background during recovery from stress induced by FUS overexpression (Figure 1E and F). These observations highlight the importance of Sbp1 in regulating the disassembly of TDP43 and FUS condensates in yeast cells.

To test the possibility that the increased accumulation of condensates in Δ*sbp1* cells was due to altered protein levels of TDP43 or FUS, we compared total protein levels using Western blotting. The relative reduction in TDP43 and FUS levels during the recovery phase was similar between wild-type and Δ*sbp1* cells (Figure 1D and G; Supplementary Figure 1B and C), indicating that the observed phenotype is not attributable to differences in protein abundance. Together, these results identify Sbp1 as a key regulator of TDP43 and FUS condensates, likely by affecting disassembly in *Saccharomyces cerevisiae*.

### TDP43 and FUS overexpression-mediated toxicity increases in Δ*sbp1*

TDP43 and FUS expression-mediated cytotoxicity have been well-documented in yeast cells^23–25,27^. Cells transformed with a galactose-inducible *TDP43/FUS*-expressing vector show condensate formation in the cytoplasm and increased cytotoxicity. To evaluate the functional relevance of Sbp1 in this model, we examined the growth of TDP43 and FUS-overexpressing cells in the Δ*sbp1* background.

Expression of TDP43 or FUS significantly impaired growth compared to empty vector controls in both wild-type and Δ*sbp1* backgrounds, as shown by spot assays (Supplementary Figure 1D and E, and Figure 1H and I). Notably, *SBP1* deletion further exacerbated the growth defect in both TDP43 and FUS-expressing cells, indicating increased sensitivity to expression in the absence of Sbp1. These findings suggest that Sbp1 counters against TDP43/FUS-mediated cytotoxicity in yeast. The enhanced toxicity observed in Δ*sbp1* cells is consistent with increased accumulation of TDP43 and FUS condensates (Figure 1B and E).

### RGG-motif protein expression reduces mutant TDP43 and FUS condensates in mammalian cells

In order to understand the role of Sbp1 as a modulator of TDP43/FUS condensates further, we examined its effect in mammalian cells. As Sbp1 lacks a known mammalian homolog, the *Saccharomyces cerevisiae SBP1* gene was cloned into mammalian expression vectors for heterologous expression. We focused on clinically relevant mutants of TDP43 and FUS that disrupt their nuclear localization and promote accumulation in cytoplasmic condensates. TDP43-ΔNLS mutant (lacking nuclear localization signal, Figure 1A) mislocalizes to the cytoplasm and forms condensates, which are reported to be toxic and clinically relevant^28–30^. TDP43-WT localized to the nucleus of HEK293T cells, whereas a significant number of cells with cytoplasmic condensates were observed when the TDP43-ΔNLS mutant was expressed (Figure 2A). However, the expression of Sbp1 significantly increased the number of cells without TDP43-ΔNLS condensates (Figure 2A and B). This phenotype was abrogated upon deletion of the RGG-motif from Sbp1 (Sbp1ΔRGG), indicating that the RGG motif is important for this activity (Figure 2A and B). Western blot analysis confirmed that the total protein levels of TDP43-WT and TDP43-ΔNLS were not significantly altered by either Sbp1 or Sbp1ΔRGG expression (Figure 2C and Supplementary Figure 2A), suggesting that the observed effects are not due to changes in protein abundance. We conclude that Sbp1 reduces the cytoplasmic condensates of TDP43-ΔNLS in mammalian cells in an RGG-motif-dependent manner.

**Figure 2:**
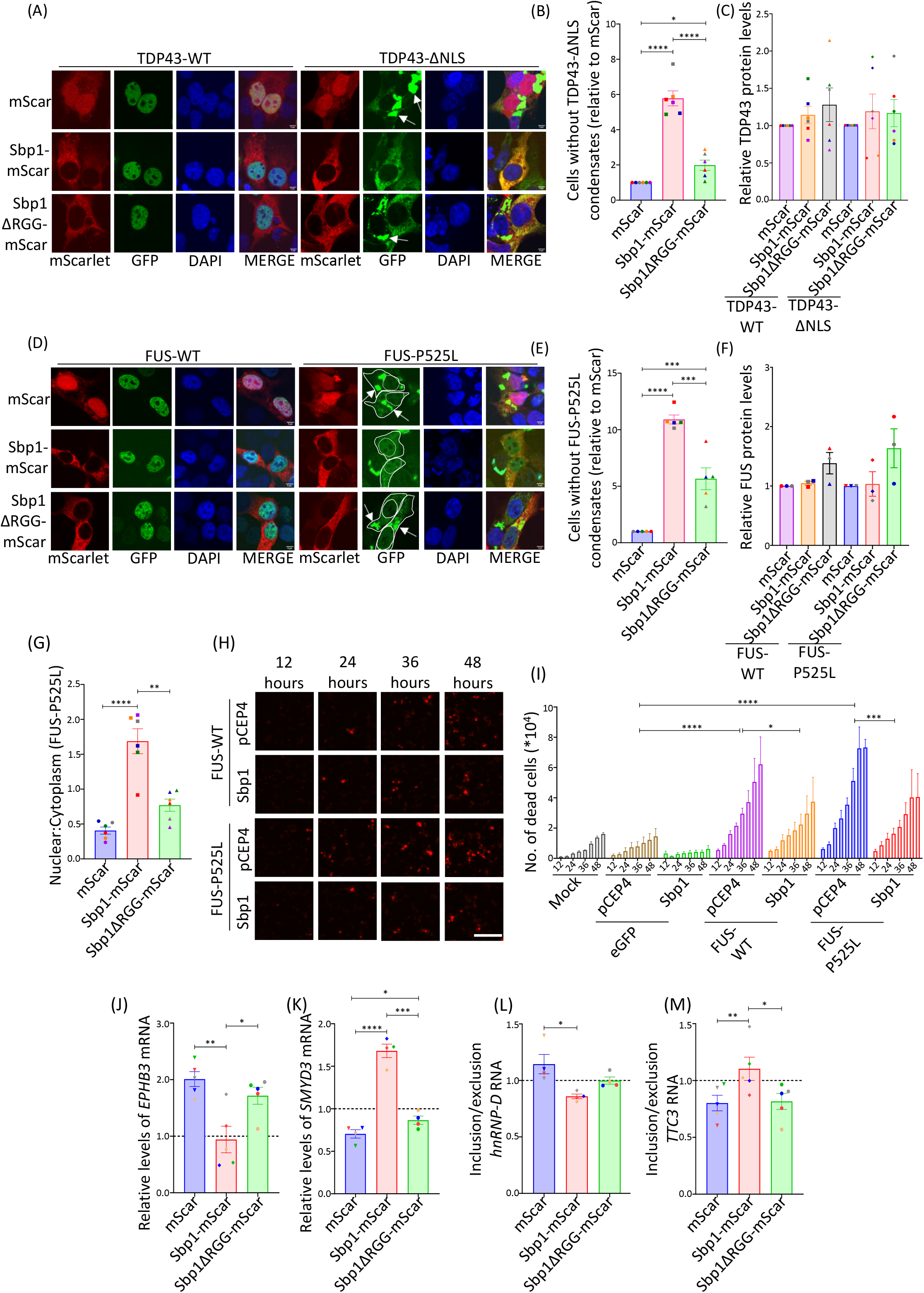
RGG-motif protein expression reduces mutant TDP43 and FUS condensates and reduces overexpression-mediated defects of FUS-P525L in mammalian cells. (A) Representative microscopy images of HEK293T cells depicting the effect of Sbp1 or Sbp1ΔRGG expression on the localisation of TDP43-WT and TDP43-ΔNLS. The white arrow marks the presence of cytoplasmic condensates. Scale bar=5um. (B) Graph representing the relative changes in the cells without TDP43-ΔNLS condensates (relative to the mScar transfected cells). The values were also normalized with the respective mScarlet fluorescent intensity expression levels. Tukey’s multiple comparisons test (2-way ANOVA) was used to calculate the significance value (n=6). (C) Graph depicting the change in TDP43-WT and TDP43-ΔNLS protein levels upon expression of mScar/Sbp1-mScar/Sbp1ΔRGG-mScar from 6 independent experiments (n=6). Tukey’s multiple comparisons test (2-way ANOVA) was used to calculate the significance value. (D) Microscopy images of HEK293T cells depicting the effect of Sbp1 or Sbp1ΔRGG expression on the localisation of FUS-WT and FUS-P525L proteins. The white arrows mark the presence of cytoplasmic condensates. Scale bar=5um. (E) Graph representing the relative changes in the cells without FUS-P525L condensates (relative to the mScar transfected cells). The values were also normalized with the respective mScarlet fluorescent intensity expression levels. Tukey’s multiple comparisons test (2-way ANOVA) was used to calculate the significance value (n=5). (F) Graph representing the change in the levels of FUS-WT and FUS-P525L in the presence of mScar/Sbp1-mScar/Sbp1ΔRGG-mScar (n=4). Tukey’s multiple comparisons test (2-way ANOVA) was used to calculate the significance value. (G) Graph representing the distribution of FUS-P525L protein in the nucleus and cytoplasm from the experiment as performed in Figure 2D. Tukey’s multiple comparisons test (2-way ANOVA) was used to calculate the significance value (n=6). (H) Incucyte images (real-time cell death analysis) representing the cellular uptake of propidium iodide (PI) in different conditions in HeLa cells. Scale bar=100um. (I) Graph representing the number of propidium iodide (PI) positive cells (dead cells) from the experiment as performed in H. The values on the x-axis reflects the time point after transfection in hours. The significance was calculated using Tukey’s multiple comparisons test (2-way ANOVA) with multiple comparisons (n=4). (J and K) Graphs representing the change in the mRNA levels of *EPHB3* (J) and *SMYD3* (K) from N2A cells expressing FUS-P525L along with either mScar/Sbp1/Sbp1ΔRGG. The values were normalised with respect to the respective eGFP-transfected conditions, denoted here by the horizontal line at 1. Tukey’s multiple comparisons test (2-way ANOVA) was used to calculate the significance values. (L and M) Graphs depicting the change in the inclusion to exclusion ratio for *hnRNP-D* (L) and *TTC3* (M) RNAs from N2A cells expressing FUS-P525L, along with either mScar/Sbp1/Sbp1ΔRGG. The values were normalised with respect to the respective eGFP-transfected conditions, denoted here by the horizontal line at 1. Tukey’s multiple comparisons test (2-way ANOVA) was used to calculate the significance values. Error bars in all graphs represent mean ±SEM, and the same color points in a graph depict the data from a single experimental set. *, **, ***, and **** denote p-value≤0.05, ≤0.01, ≤0.001, and ≤0.0001, respectively.

A similar experiment was conducted for FUS-WT and the P525L mutant. This mutation is in the NLS motif of the protein and has been correlated with an aggressive form of juvenile ALS^31^ (Figure 1A). Consistent with previous reports, expression of FUS-P525L in HEK293T cells led to robust cytoplasmic condensate formation, while FUS-WT localized predominantly to the nucleus^32–34^ (Figure 2D). Co-expression of Sbp1 significantly increased the proportion of cells lacking FUS-P525L condensates (Figure 2D and E). This effect was markedly diminished with the Sbp1ΔRGG variant, mirroring results observed with TDP43 (Figure 2D and E). This phenotype was not due to a change in FUS protein levels, as the Western analysis reflected no significant change in FUS protein in any of the conditions (Figure 2F and Supplementary Figure 2B). Together, these data demonstrate that yeast-derived Sbp1 can reduce cytoplasmic condensates formed by ALS-associated TDP43 and FUS mutants in mammalian cells. This activity requires the RGG-motif of Sbp1 and occurs independently of changes in total protein levels.

### RGG-motif protein expression promotes nuclear re-localization of FUS-P525L and reduces overexpression-mediated toxicity

In addition to reducing cytoplasmic P525L condensate, we investigated whether Sbp1 could restore nuclear localization of the ALS-linked FUS-P525L mutant in mammalian cells. Quantification of the nuclear-to-cytoplasmic fluorescence ratio revealed that Sbp1 expression led to an increase in the nuclear localization of the mutant FUS (Figure 2G). While the RGG-motif deletion mutant was able to rescue cells without condensates to a lesser extent, it was significantly defective in rescuing the nuclear localization defect as compared to the wild-type Sbp1 (Figure 2G). No change in the localization of FUS-WT was observed in the presence of either Sbp1 or Sbp1ΔRGG.

FUS overexpression is associated with increased cytotoxicity^35^; therefore, we asked whether Sbp1 could mitigate this phenotype. Cells transfected with FUS WT or FUS-P525L exhibited a marked increase in cell death, as measured by propidium iodide (PI) staining over time using an Incucyte live-cell imaging system (Figure 2H and I, and Supplementary Figure 2C). While not significant, we also observed slightly increased toxicity of FUS-P525L at all time points of our analysis. We observed a significant reduction in the toxicity of both FUS-WT and P525L in the presence of Sbp1; however, Sbp1 did not significantly affect the cell viability of eGFP-transfected cells (Figure 2H and I). These results suggest that Sbp1 not only reduces mutant FUS condensate burden but also facilitates its nuclear relocalization and suppresses FUS-mediated cytotoxicity in mammalian cells.

### RGG-motif protein expression rescues FUS-P525L-induced defects in RNA abundance and splicing

FUS is a multifunctional RNA-binding protein involved in regulating transcript stability and alternative splicing, and its loss or mutation leads to widespread RNA misregulation^35,36^. To assess whether Sbp1 can restore the functional activity of FUS, we examined known FUS targets affected by its mislocalization or knockdown^35,36^. Specifically, we analyzed the expression and splicing of four FUS-regulated genes in mouse neuroblastoma cells: *Ephb3* and *Smyd3*, which are regulated at the level of RNA stability, and *hnRNP-D* and *Ttc3*, which are regulated through alternative splicing in various mouse models^35,36^.

Consistent with previous studies in mouse models (with either expression of FUS-ΔNLS or FUS knockdown), overexpression of FUS-P525L led to upregulation of *Ephb3* and downregulation of *Smyd3*, reflecting impaired FUS function (Figure 2J and K). Expression of Sbp1 significantly rescued the aberrant expression of both transcripts, restoring them toward wild-type levels (Figure 2J and K). This rescue was dependent on the RGG-motif, as the Sbp1ΔRGG mutant failed to rescue the transcript levels. Similarly, FUS-P525L overexpression resulted in splicing defects in *hnRNP-D* (increased exon inclusion) and *Ttc3* (decreased exon inclusion) (Figure 2L and M, and Supplementary Figure 2D and E). Sbp1 expression restored normal splicing of both targets, again in an RGG-motif-dependent manner (Figure 2L and M, and Supplementary Figure 2D and E). These results indicate the role of Sbp1 in rescuing the P525L-associated functional defect.

### Sbp1 directly binds FUS and disassembles condensates in an RGG-motif-dependent manner

The observed reduction in cytoplasmic condensates upon Sbp1 expression could result from impaired assembly or enhanced disassembly of pre-formed aggregates. To explore this further, we first tested whether Sbp1 physically interacts with FUS. Co-immunoprecipitation experiments using mScarlet-tagged Sbp1 revealed that Sbp1 binds FUS-P525L, and this interaction was highly compromised upon deletion of the RGG-motif (Sbp1ΔRGG), indicating that the RGG region is essential for binding to FUS (Figure 3A and B). We further tested Sbp1 and FUS binding using purified proteins (all purified protein used in this study are shown in Supplementaty Figure 3A). Full length Sbp1 interacts with FUS however the RGG-deletion mutant was significantly defective in binding to FUS (Figure 3C).

**Figure 3:**
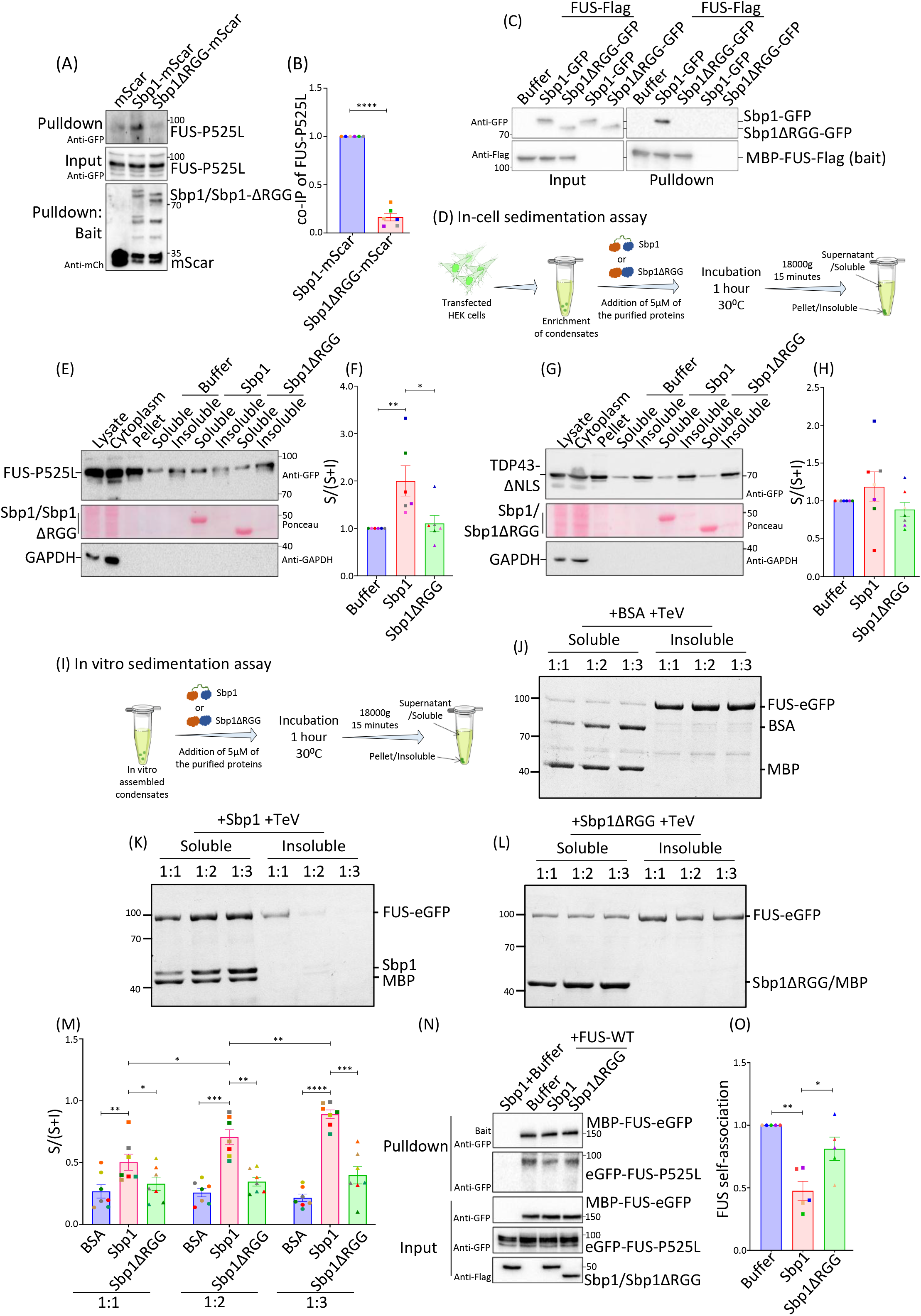
RGG-motif protein directly disassembles FUS condensates by disrupting FUS self-association through an RGG-motif-dependent mechanism. (A) Western analysis for the experiement with pulldown of Sbp1/Sbp1ΔRGG (bait: mScarlet tag) depicting the interaction of Sbp1 with FUS-P525L. Values on the right represent the position of different molecular weight ladder bands in kDa. (B) Quantitation for the amount of FUS-P525L in the pulldown fraction from the experiment as performed in A. The values were normalised with respect to the amount of bait in the pulldown reaction. A student-paired t-test analysis calculated significance (n=6). (C) Western analysis depicting the pulldown of Sbp1/Sbp1ΔRGG with FUS-Flag (bait) from an *in vitro* pulldown experiment. Values on the right represent the position of different molecular weight ladder bands in kDa. (D) Schematic depicting the workflow for the in-cell sedimentation assay used to assess the disassembly activity of Sbp1. (E and G) Western analysis of the different fractions from FUS-P525L (E), and TDP43-ΔNLS (G) in-cell sedimentation assay. The different fraction loadings are as follows: lysate: 7.5%, cytoplasm: 7.5%, pellet: 15%, soluble: 60%, and insoluble: 60%. GAPDH serves as the control for the assay, and ponceau reflects the purified protein added to the respective reaction. Values on the right represent the position of different molecular weight ladder bands in kDa. (F and H) Quantitation of the fraction of protein in the soluble fraction from experiments as done in E and G. Significance was calculated by using Tukey’s multiple comparisons test (2-way ANOVA; n=6 and n=7 for E and G, respectively). (I) Schematic depicting the workflow of the *in vitro* sedimentation assay performed to assess the disassembly activity of Sbp1 on the phase-separated FUS and TDP43 condensates. (J-L) Coomassie-stained protein gels depicting the fractionation of FUS to soluble and insoluble phases in the presence of BSA (J), Sbp1 (K), and Sbp1ΔRGG (L). The ratio reflects the amount of FUS: test protein taken for the assay. Sbp1ΔRGG and MBP migrate at the same position and hence appear as a single band. Values on the left represent the position of different molecular weight ladder bands in kDa. (M) Quantitation of the fraction of FUS protein present in the soluble phase from 7 independent experiments (n=7) as performed in J-L. Significance was calculated by Tukey’s multiple comparisons test (2-way ANOVA). (N) Western analysis depicting the interaction of eGFP-FUS-P525L upon pulldown of MBP-FUS-eGFP in the presence of Sbp1/Sbp1ΔRGG. Sbp1+Buffer lane had everything except the bait, MBP-FUS-eGFP. (O) Quantitation for the amount of FUS-P525L from the pulldown experiment as performed in N. The values were also normalised with respect to the amount of bait in the pulldown reaction. Tukey’s multiple comparisons test (2-way ANOVA) was used to calculate the significance (n=5). Error bars represent mean ±SEM, and the same color points depict the data from a single experimental set. *, **, ***, and **** denote p-value ≤0.05, ≤0.01, ≤0.001, and ≤0.0001, respectively.

Further, we directly assessed the disassembly activity of Sbp1 on mutant FUS and TDP43 condensates from mammalian cells by using a modified in-cell sedimentation assay^37^. Recombinant Sbp1 protein was purified (Supplementary Figure 3A) and incubated with the enriched condensates at 30^0^C for 1 hour, followed by differential centrifugation to separate soluble (supernatant) and insoluble (pellet) fractions (Figure 3D and see methods). Western blot analysis revealed that in the control (buffer-only) condition, FUS-P525L was predominantly retained in the pellet fraction, consistent with its localisation in cytoplasmic condensates (Figure 3E). In contrast, Sbp1 treatment led to a substantial redistribution of FUS-P525L into the soluble fraction (Figure 3E and F), indicating disassembly. This shift was not observed with the Sbp1ΔRGG mutant, which showed significantly impaired activity (Figure 3E and F). As expected, GAPDH, a soluble cytoplasmic protein, was exclusively present in the supernatant in all conditions (Figure 3E and G). Notably, Sbp1 had no measurable disassembly effect on TDP43-ΔNLS condensates in this assay (Figure 3G and H), suggesting a distinct underlying mechanism for TDP43 disassembly.

To determine whether Sbp1 could directly act on condensates in the absence of other cellular components, we performed an *in vitro* two-component sedimentation assay using purified recombinant FUS and TDP43 proteins (Figure 3I). Purified proteins used in these experiments are shown in Supplementary Figure 3A. Following *in vitro* phase separation to form condensates (Supplementary Figure 3B and C), the proteins (phase-separated) were incubated with either Sbp1, Sbp1ΔRGG, or BSA as a control. Incubation with Sbp1 resulted in a concentration-dependent increase in the soluble fraction of FUS, indicative of condensate disassembly (Figure 3K and M). This effect was not observed in BSA-treated controls, and Sbp1ΔRGG was markedly less effective in promoting solubilization (Figure 3J, L, and M). To test the importance of arginine residues and/or arginine methylation within the RGG-motif, we generated an RGG-mutant variant (Sbp1AMD) in which all arginine residues were substituted with alanine, rendering the motif defective in arginine methylation. Like the RGG deletion mutant, Sbp1AMD failed to disassemble FUS condensates (Supplementary Figure 3D and E), highlighting the critical role of arginine-mediated interactions in the disassembly mechanism.

### FUS self-association is disrupted by Sbp1 in RGG-motif dependent manner

To dissect the molecular mechanism by which Sbp1 disassembles FUS condensates, we next examined whether it could interfere with FUS self-association. Recombinant MBP-FUS-eGFP protein was incubated with lysates from HEK293T cells expressing FUS-P525L, and MBP pull-down was performed to detect FUS-FUS interactions. In the absence of Sbp1, FUS-P525L co-purified with MBP-FUS-eGFP, confirming self-association (Figure 3N and O).

Addition of recombinant Sbp1 significantly reduced this interaction, indicating competitive disruption of FUS-FUS binding. This disruption was significantly reduced when the RGG-motif was deleted from Sbp1, confirming that the RGG region is essential for its ability to outcompete FUS self-association (Figure 3N and O). The direct nature of competition was also confirmed using recombinant purified proteins (Supplementray Figure 3F and G). Together, these results identify a direct, RGG-motif-dependent role for Sbp1 in disassembling FUS condensates by engaging FUS and competitively interfering with its self-interactions.

### RGG-motif is sufficient to disassemble ALS-relevant condensates

Previous experiments established the important role of the RGG-motif in the disassembly of mutant FUS condensates. To test whether the RGG-motif is sufficient for this activity, we first examined the disassembly potential of different peptides by using an in-cell sedimentation assay. Three peptides were tested: (i) the disordered RGG-motif (residues 123 to 170; 48 amino acids) of Sbp1 (Sbp1-RGG), (ii) the disordered RGG-motif (residues 283 to 331; 49 amino acids) of another yeast protein, Scd6 (Scd6-RGG), and (iii) a peptide from the RRM domain (residues 37 to 85; 49 amino acids) of Sbp1 (Sbp1-RRM) (Figure 4A). The disordered nature of the peptides was checked by AIUPred analysis (Supplementary Figure 4A). Among the three, only the Sbp1-RGG peptide effectively redistributed FUS-P525L from the insoluble to the soluble fraction, similar to the full-length Sbp1 (Figure 4B and C). Neither Scd6-RGG nor Sbp1-RRM induced any measurable change in FUS solubility, suggesting a sequence and context-specific disassembly function encoded within the Sbp1-RGG motif.

**Figure 4:**
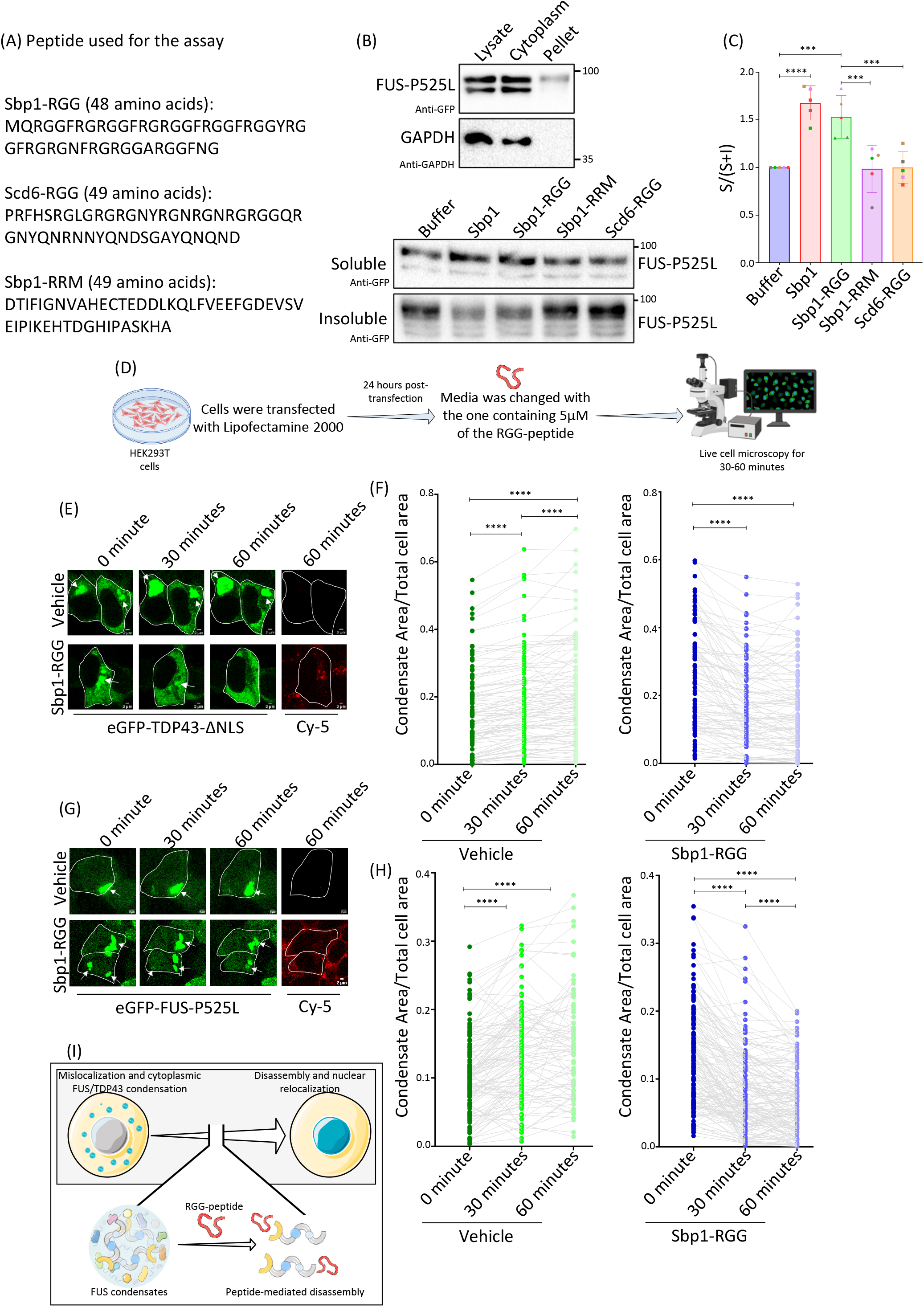
RGG peptide disassembles the TDP43-ΔNLS and FUS-P525L condensates. (A) The sequence of the peptides used for the experiments. For Sbp1-RGG, 2 extra amino acids at the N-terminal end (M and Q) and 3 amino acids at the C-terminal end (F, N, and G) were included for better purification of the RGG-peptide. (B) Western analysis of the different fractions from FUS-P525L in-cell sedimentation assay using various peptides. The different fraction loadings are as follows: lysate: 7.5%, cytoplasm: 7.5%, pellet: 15%, soluble: 60%, and insoluble: 60%. GAPDH serves as the control for the assay. Values on the right represent the position of different molecular weight ladder bands in kDa. (C) Quantitation of the fraction of protein in the soluble fraction from experiments as done in B. Significance was calculated by using Tukey’s multiple comparisons test (2-way ANOVA, n=5). (D) Schematic depicting the workflow of live-cell microscopy with the RGG-peptides. (E and G) Microscopy images for HEK293T cells depicting the change in the area of TDP43-ΔNLS (E) and FUS-P525L (G) condensate (marked by white arrows) after incubating with either the vehicle control or Sbp1-RGG peptide. Cy5 depicts the peptide uptake after 60 minutes of incubation with the peptide. Scale bar=2um. (F and H) Graphs depicting the quantitation of the condensate area/total cell area from at least 30 cells per experiment (n=3). Significance was calculated by paired t-test analysis. (I) Schematic depicting the role of RGG peptides as a therapeutic approach to disassemble the toxic cytoplasmic condensates. Blue represents mutant proteins, like FUS-P525L and TDP43-ΔNLS, that mislocalize to the cytoplasm and form condensates in diseased conditions. RGG peptides could have the potential to disassemble these toxic condensates. The inset also depicts the possible mechanism of the disassembly of FUS condensates by RGG-peptide. Disassembly may also result in the restoration of the nuclear localization phenotype. Error bars represent mean ±SEM, and the same color points depict the data from a single experimental set. *** and **** denote p-value ≤0.001 and 0.0001, respectively.

To assess whether the Sbp1-RGG peptide is sufficient to disassemble mutant condensates in mammalian cells, we performed live-cell imaging following exogenous peptide addition. Sbp1-RGG peptide was added to the culture media of cells expressing either FUS-P525L or TDP43-ΔNLS, followed by live-cell microscopy (Figure 4D). While both FUS and TDP43 condensate size increased for the vehicle control, there was a significant reduction in size within 30 minutes of peptide treatment (Figure 4E–H). Continued incubation for an additional 30 minutes led to a further decrease in condensate size, consistent with progressive disassembly.

Together, these results demonstrate that the RGG-motif is sufficient to trigger the disassembly of pathological condensates formed by ALS-associated FUS and TDP43 mutants. These findings identify Sbp1-RGG as a minimal disassembly-competent unit and highlight its potential as a peptide-based therapeutic strategy for targeting aberrant RNP condensates in ALS.

## Discussion

Aberrant cytoplasmic condensation of nuclear RNA-binding proteins, particularly FUS and TDP43, is a hallmark of ALS and related neurodegenerative disorders. While substantial effort has focused on understanding condensate assembly and clearance, the disassembly of these pathological assemblies remains poorly characterized. In this study, we identify the intrinsically disordered RGG-motif of the yeast protein Sbp1 as both necessary and sufficient to disassemble ALS-relevant FUS-P525L and TDP43-ΔNLS condensates *in vivo*. Through a combination of yeast genetics, mammalian cell biology, and biochemical reconstitution, we establish the RGG-motif as a bona fide disassembly-inducing peptide and uncover a novel mechanism to mitigate condensate-associated toxicity.

*Saccharomyces cerevisiae* has long served as a genetically tractable model to study protein aggregation in neurodegenerative diseases, including ALS^14–19^. Prior screens have identified numerous modifiers of FUS^20,26^ and TDP43^13^ toxicity in yeast, yet none, to our knowledge, have explored the identification of disassembly-promoting factors. Motivated by prior evidence implicating the RGG-motif of Sbp1 in processing body (PB) disassembly^38^, we hypothesized that Sbp1 could disassemble TDP43 and FUS condensates. Interestingly, TDP43 and FUS also have long stretches of LCS region, with FUS having N-terminal QGSY and an RGG-motif, and TDP43 having a long C-terminal glycine-rich region (Figure 1A). Indeed, using a galactose-inducible overexpression system, we observed a defect in the disassembly, in Δ*sbp1* cells, accompanied by enhanced cytotoxicity. This effect was not due to altered protein levels, suggesting a direct role of Sbp1 in regulating phase-separated assemblies. Our results are supported by the observation that overexpression of Sbp1 suppresses FUS overexpression toxicity in yeast cells^20,26^. However, these studies did not assess the underlying mechanism of the regulation of toxicity by Sbp1. Importantly, our study characterizes the crucial role of the disordered domain of Sbp1 in disassembling ALS-related condensates, a process that was previously unexplored. No such connection has been identified for TDP43 so far. Therefore, our observations establish Sbp1 as a novel and specific regulator of mutant TDP43 and FUS condensate disassembly in yeast.

Based on our results with yeast cells, we were motivated to check the effect of Sbp1 expression on TDP43 and FUS condensates in mammalian cell models. The cytoplasmic mislocalization and aggregate formation are hallmark features of TDP43 in both familial and sporadic ALS cases^39^. Similarly, FUS mislocalization and aggregate formation are well-reported in familial cases^40^. Different mutations have been identified in both TDP43 and FUS that can enhance the rate of these defects. Overexpression of such mutant forms recapitulates the ALS-related phenotype and has been instrumental in understanding different aspects of the disease^41,42^. In our experiments, we expressed TDP43-ΔNLS and FUS-P525L mutants that mislocalized and formed cytoplasmic condensates (Figure 2). Interestingly, Sbp1 expression significantly increased the number of cells without cytoplasmic condensates of the mutant forms of TDP43 and FUS. Importantly, this activity was dependent on the RGG-motif, though modest residual activity in the RGG-deletion mutant suggests that the RNA recognition motifs (RRMs) may also contribute towards regulating the condensate dynamics.

The toxic phenotype of the mutant FUS has been associated with both its cytoplasmic mislocalization and condensate formation^33,42^. In addition to reducing cytoplasmic FUS-P525L condensates, Sbp1 promoted its nuclear localization and significantly enhanced cell viability (Figure 2G-I). Furthermore, Sbp1 rescued the expression and splicing defects of several FUS-regulated targets, including *Ephb3* and *Smyd3* (whose stability is affected, Figure 2J and K), and *hnRNP-D* and *Ttc3* (whose splicing is affected, Figure 2L and M). It is noteworthy that the RGG-motif deletion mutant was unable to induce FUS relocalization and rescue the defects associated with its RNA targets. Such an observation highlights the essential role of the RGG-motif in the disassembly and rescue of the functional defects of FUS-P525L in these conditions. These observations open an altogether new direction for exploring the role of disordered sequences as disassembly factors.

Our experiments further revealed mechanistic insights into the role of the RGG-motif in modulating cytoplasmic condensates of mutant FUS and TDP43. We observed a direct interaction between Sbp1 and FUS (Figure 3A-C), which was dependent on the Sbp1 RGG-motif, highlighting the ability of the disordered RGG sequence to target FUS. Using both modified in-cell and *in vitro* sedimentation assays, we demonstrated that wild-type Sbp1, but not its RGG-deletion or arginine-substitution mutant, could disassemble pre-formed FUS-P525L condensates (Figure 3E, K and L, and Supplementary Figure 3D and E). These findings underscore the critical role of arginine-mediated multivalent interactions, such as cation-π and π-π stacking, in driving condensate disassembly. Notably, FUS itself relies on its low-complexity sequences, including RGG motifs (interaction between arginine of the RGG-motif and tyrosine of the QGSY-region), for phase separation^43–46^, raising the possibility that the RGG-motif of Sbp1 engages in competitive interactions with those of FUS. Supporting this, co-immunoprecipitation experiments from cells and direct interaction experiments using recombinant purified proteins revealed an RGG-motif-dependent (of Sbp1) interaction between Sbp1 and FUS (Figure 3A-C). Furthermore, Sbp1 effectively disrupted FUS: FUS self-association both in cell lysate and *in vitro*, and this disruption was abolished upon deletion of the RGG-motif of Sbp1 (Figure 3N and O, and Supplementary Figure 3F and G). These results suggest that Sbp1 mediates condensate disassembly by directly engaging FUS and competitively interfering with its self-association, thereby promoting the disassembly of pre-formed assemblies.

Interestingly, while Sbp1 robustly disassembled FUS condensates, it failed to disassemble enriched and pre-formed TDP43-ΔNLS condensates, despite clear disassembly activity in cells (Figure 3G and H). This discrepancy suggests that disassembly of TDP43 condensates may rely on additional cofactors or a more complex cellular context. Alternatively, differences in the material properties or molecular architecture of FUS and TDP43 condensates may affect their susceptibility to Sbp1-mediated disassembly. This observation highlights that condensate-specific disassembly factors and mechanisms are likely to exist, and exploring these should be an important avenue for researchers in the field. Understanding the molecular basis underlying the lack of sensitivity of the mutant TDP43 condensates to purified Sbp1 would be a future endeavour for our group. A recent report identified the proteome of the insoluble TDP43 fraction from the brain tissue of TDP43-ΔNLS mice^47^. Identifying some intermediate players from this study that can be further directed to induce the disassembly of toxic TDP43 condensates will be an interesting direction. However, the sensitivity of both FUS-P525L and TDP43-ΔNLS to RGG peptides in mammalian cells is encouraging to further explore possible therapeutic applications of RGG peptides in ALS.

Different kinds of condensate targeting molecules have been identified, and many of these primarily target the assembly of proteins into the condensates^5–9^. The list includes many small molecules and a few peptides. Small planar compounds, like mitoxantrone, have been identified as affecting both the assembly and disassembly of mutant TDP43 condensates, reducing the cumulative death rate of the primary neurons^48^. Apart from these, the current literature on peptides targeting TDP43 only reports the degradation-promoting peptides^49,50^. These peptides were designed to have a TDP43 recognition motif, which is a part of the TDP43 protein that has the ability to self-associate. No such peptides are reported for FUS condensates to the best of our knowledge. Our report, for the first time, identifies a disordered RGG peptide that functions as a genuine disassembly factor (Figure 4). We demonstrate that Sbp1-RGG peptides are sufficient to disassemble both FUS-P525L and TDP43-ΔNLS condensates *in vivo*. Importantly, this effect was specific to the Sbp1-RGG motif, as neither a related RGG motif from Scd6 nor a control peptide from the RRM domain of Sbp1 showed any activity in the disassembly assay. Although condensates were disassembled by Sbp1-RGG, technical limitations prevented us from assessing whether FUS-P525L relocalized to the nucleus under these conditions. Nevertheless, the impaired nuclear relocalization observed upon expression of Sbp1ΔRGG points toward an RGG motif– dependent mechanism (Figure 2G). Further experiments will be necessary to determine whether this dependency extends to the RGG peptide.

Such insight has opened a new direction for exploring the role of disordered peptides as the disassembly factors of other disease-relevant RNP condensates. Our study provides proof of principle to explore the role of RGG-peptides as a therapeutic avenue for treating ALS (Figure 4I). Furthermore, unlike antisense oligonucleotides (ASOs) or targeted degradation approaches, which eliminate the protein entirely and may compromise nuclear functions (leading to potential loss-of-function effects), disassembly factors like the RGG peptide reported in this work offer a mechanistic advantage. They not only reduce toxic cytoplasmic condensates but also promote nuclear relocalization, as seen with FUS-P525L (Figure 2G). Therefore, a therapeutic option that specifically targets the disassembly of TDP43/FUS condensates could, in principle, also rescue the nuclear functions. Future work will be required to test RGG-peptide efficacy in disease-relevant models such as patient-derived motor neurons and to uncover the precise molecular partners that mediate TDP43 disassembly. It will also be important to examine whether such peptides influence other phase-separated condensates, as their activity is likely to be context-dependent. Such specificity would provide a valuable tool for selectively modulating the material properties of distinct condensates, similar to what has recently been demonstrated for a certain micropeptide^51^. Nonetheless, our results establish disordered RGG-peptides as functional disassembly factors and set the stage for developing targeted interventions to reverse condensate pathology in ALS.

## Materials and methods

### Yeast strain and growth conditions

The yeast strains used in this study are listed in Table 1. Strains were grown at 30^0^C in yeast extract-peptone (YP) medium, and cells with *FUS* and *TDP43* overexpressing plasmids were maintained in synthetic defined (SD) uracil dropout media (SD-Ura^-^) medium supplemented with 2% raffinose. For secondary cultures, cells were diluted to OD_600_ 0.1 and grown till the mid-log phase of OD_600_ 0.4-0.5. For protein induction, the cells were shifted to 2% galactose-containing media for the mentioned time after the mid-log phase. Recovery experiments were carried out in 2% glucose-containing SD-Ura^-^ media.

**Table 1:**
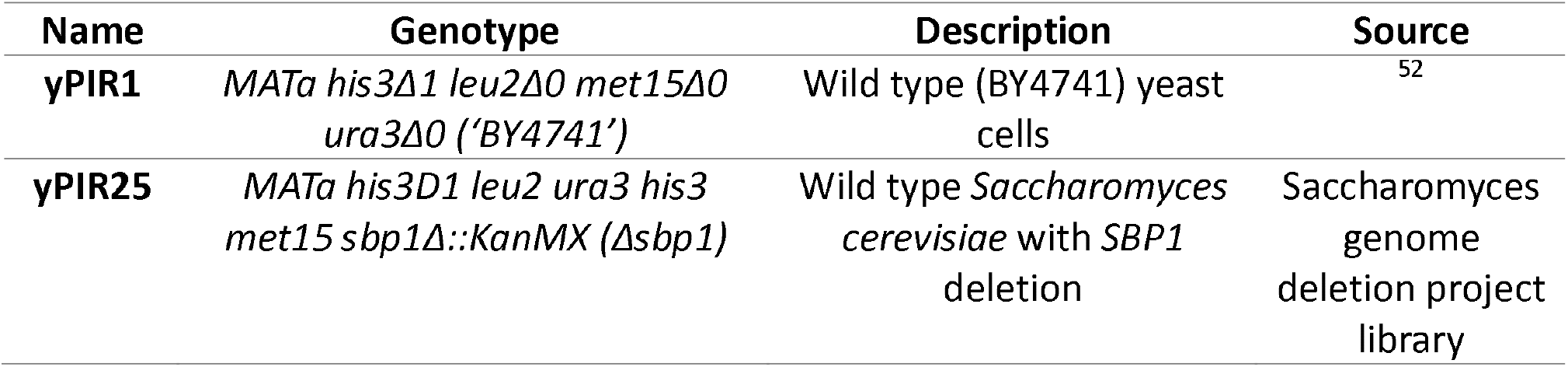
List of yeast strains used in this study.

### Yeast spot assays

Cells were grown till the mid-log phase in SD-Ura^-^ media supplemented with raffinose. *TDP43* and FUS induction were carried out by shifting the cells to 2% galactose-containing media for 2 and 3 hours, respectively. Post-induction cells were further processed for spotting assays. For spotting assays, cells were serially diluted from 10.0 OD_600_ to 0.001 OD_600_ and spotted on the SD-Ura^-^ glucose and galactose-containing plates. After sufficient growth, the images were acquired, and the spot area was analyzed as described earlier in Petropavlovskiy et al. 2020^53^.

### Plasmids

The list of plasmids used in this study is listed in Table 2. pCEP4*-HIS-SBP1-FLAG* was constructed by amplifying *HIS-SBP1-FLAG* ORF from pPROEx-*HIS-SBP1-FLAG* construct. The primers (Table 3) were designed using the NEBuilder primer design tool to keep the His and Flag tags intact at the N and C-terminus, respectively, and target the amplicon to the BamHI digested pCEP4 plasmid. Positive clones were confirmed using a PCR reaction, and the expression was checked by transfecting the plasmid into HEK293T cells, followed by Western blotting. A similar cloning strategy was also used to clone *pcDNA-SBP1*/Sbp1ΔRGG-*mScarlet*, which was cloned into the BamH1-digested pcDNA-mScarlet vector.

**Table 2:** List of plasmids used in this study.

| Name | Description | Source |
| --- | --- | --- |
| pPIR92 | pRS316 Empty vector, with *URA3* and ampicillin resistance genes | This study |
| pPIR74 | Plasmid expressing *hFUS-YFP* under a galactose inducible promoter, with *URA3* and ampicillin resistance genes | Addgene |
| pPIR75 | Plasmid expressing *hTDP43-GFP* under a galactose inducible promoter, with *URA3* and ampicillin resistance genes | Addgene |
| **pPIR29** | *E. coli* expression vector for Sbp1 with N-terminal His and C-terminal Flag tag, Ampicillin resistance | 54 |
| **pPIR33** | *pPIR29* with amino acid 125-167 deleted from the Sbp1-ORF; Sbp1ΔRGG | 54 |
| **pPIR34** | *pPIR29* with Sbp1R125A, R129A, R131A, R135A, R137A, R141A, R145A, R149A, R153A, R155A, R159A, R161A and R165A; Sbp1-AMD | 54 |
| **pPIR317** | pCEP4, vector with CMV promoter, hygromycin, and ampicillin resistance genes | Kind gift from Prof. K. Somasundaram, IISc |
| **pPIR337** | pCEP4-His-Sbp1-Flag, pCEP4 expressing N-terminal His and C-terminal Flag-tagged *SBP1*, CMV promoter | This study |
| **pPIR308** | peGFP-C1, the vector expressing eGFP under a CMV promoter | Kind gift from Prof. Sandeep M Eswarappa, IISc |
| **pPIR343** | peGFP-FUS-WT, the vector expressing eGFP-hFUS-WT under a CMV promoter | Kind gift from Dr. Dorothee Dormann, IMB |
| **pPIR344** | peGFP-FUS-P525L, the vector expressing eGFP-hFUS-P525L mutant under a CMV promoter | Kind gift from Dr. Dorothee Dormann, IMB |
| **pPIR313** | pDEST eGFP only, the vector expressing eGFP under a hybrid CMV and doxycycline-inducible promoter | 30 |
| **pPIR314** | pDEST TDP43-WT, pPIR313 with hTDP43 cloned upstream of eGFP | 30 |
| **pPIR315** | pDEST TDP43-ΔNLS, pPIR313 with hTDP43-ΔNLS cloned upstream of eGFP | 30 |
| **pPIR365** | pcDNA-mScarlet empty vector | Kind gift from Dr. Janin Lautenschlager |
| **pPIR364** | pcDNA-Sbp1-mScarlet | This study |
| **pPIR373** | *pPIR364* with amino acids 125-167 (from the Sbp1-ORF which starts at 22<sup>nd</sup> amino acid) deleted; Sbp1ΔRGG-mScarlet | This study |
| **pPIR375** | *E. coli* expression vector pMAL-MBP-TeV-FUS-eGFP-TeV-His | Kind gift from Dr. Dorothee Dormann, IMB<sup>43</sup> |
| **pPIR377** | *E. coli* expression vector, expressing His-TEV in a pET-24d(+) vector | Kind gift from Dr. Dorothee Dormann, IMB<sup>43</sup> |
| **pPIR393** | *E. coli* expression vector pMAL-MBP-TeV-FUS-Flag-TeV-His | Kind gift from Dr. Dorothee Dormann, IMB<sup>43</sup> |
| **pPIR57** | *E. coli* expression vector pPROEx-His-Sbp1-GFP | This paper |
| **pPIR58** | *E. coli* expression vector pPROEx-His-Sbp1-Sbp1ΔRGG-GFP | This paper |

**Table 3:**
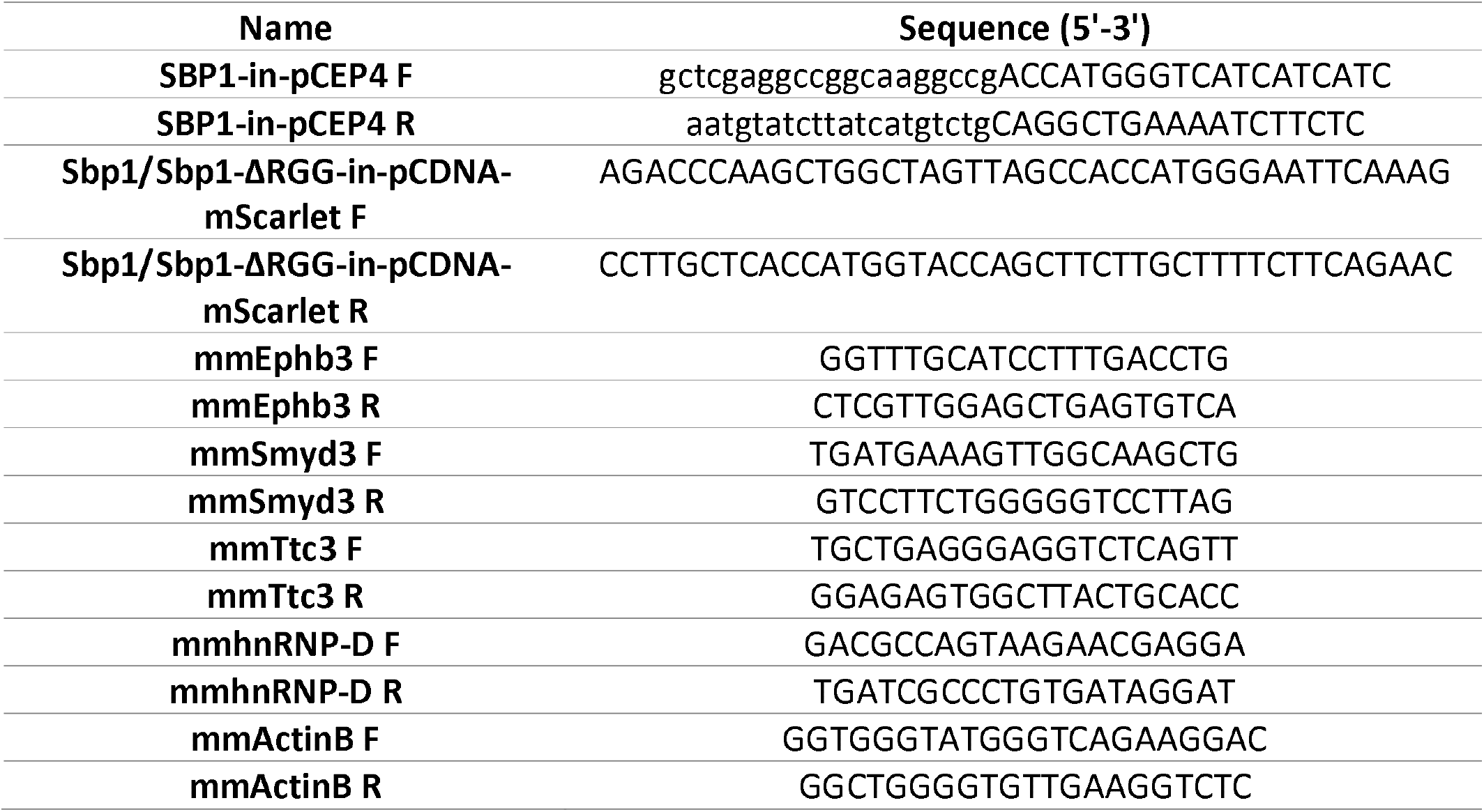
List of primers used in this study.

### Mammalian cell cultures

HEK293T and HeLa cells were maintained in Dulbecco’s Modified Eagle Medium (DMEM) supplemented with 10% FBS and 1X antibacterial-antimycotic solution (complete DMEM). Media was changed every 24 hours for proper growth and split after ≥90% confluency. Cultures were checked for Mycoplasma contamination once every month by a PCR-based method.

### Transfection and preparation of mammalian cell samples for microscopy analysis

For transfection, Lipofectamine 2000 or 3000 (Thermo) was used as per the manufacturer’s protocol. The cells were grown, transfected, and processed either directly on culture plate (for Western and RNA analysis) or coverslips (for microscopy). The samples were collected 24 hours post-transfection, and the cells were fixed with 4% formaldehyde solution for 15-20 minutes. Three washes were given with 1X PBS. For the experiment in Figure 2A and D, the coverslips were directly mounted onto slides with Fluoromount-G containing DAPI and stored at 4^0^C until imaging was done.

For the live-cell peptide uptake assay, cells were grown and transfected on 35mm glass-bottom dishes, and the plate was directly taken for microscopy. Images were acquired at 63X objective in an incubation chamber with 5% CO_2_ at 37^0^C. After fixing the fields, the media was changed to a fresh one containing 5μM of the Cy5-labeled peptides (synthesized from Genscript). The imaging was done for 60 minutes, with images taken at every 15-minute interval. After 60 minutes, the cells were washed in PBS thrice and resuspended in PBS to check the uptake of Cy5-labeled peptides.

### Mammalian cell viability assay

The cell viability assay was performed in the Incucyte chamber. Briefly, after 6 hours of transfections, the cell medium was supplemented with 500uM propidium iodide (PI), and images were acquired every hour to score the number of PI-positive cells. The cell death analysis was performed using the IncuCyte S3 live-cell analysis instrument (Sartorius), and the change in the number of PI-positive cells (dead cells) in different conditions was plotted in the graph.

### RNA abundance and splicing assays

The total RNA from N2A cells was isolated using Trizol. Briefly, pelleted cells were resuspended in 100ul of PBS. 1ml trizol and 200ul of chloroform were added to the cells and vortexed for 15-30 seconds. The solution was allowed to rest at 15-30^0^C for 3 minutes. The aqueous phase was separated by centrifugation at 12000g for 15 minutes at 4^0^C. After centrifugation, the aqueous phase was collected in a fresh tube, and 500ul of isopropanol was added. The reaction was allowed to incubate at 15-30^0^C for 10 minutes, and the RNA was precipitated at 12000g for 15 minutes at 4^0^C. This was followed by two washes in 70% ethanol with centrifugation at 7500g for 5 minutes each. The final pellet was allowed to air dry and resuspended in 30ul of nuclease-free water. The quality and quantity of the RNA were checked by 1.2% agarose formamide gel electrophoresis and nanodrop, respectively. Four microgram of RNA was used to synthesize cDNA using the RevertAid reverse transcriptase as per the manufacturer’s protocol. cDNA was diluted to 1/10, and PCRs were carried out with the respective exon-specific primer sets (Table 3).

Real-time PCR with SYBR green dye was used for the measurement of RNA abundance. For this, two technical replicates were assembled with 2ul cDNA/reaction and 0.5uM each primer in a BioRad iQ5 Real-Time PCR Detection System. The PCR conditions were 95^0^C for 10 minutes for initial denaturation, followed by 35 cycles of 95^0^C for 15 seconds, and 60^0^C for 45 seconds. DNA was quantified in every cycle at the extension step. Melt curve acquisition was carried out from 60^0^C to 95^0^C with an increment of 0.5^0^C. Ct values were extracted with auto baseline and manual threshold. ^ΔΔ^Ct method was used to calculate the final log_2_-FoldChange values and plotted using GraphPad Prism 8.0. ActinB served as the internal control.

For splicing assays, a regular PCR reaction was set up by using 2ul cDNA in a 25ul reaction. The PCR conditions were 98^0^C for 5 minutes for initial denaturation, followed by 35 cycles of 98^0^C for 1 minute, 50^0^C for 45 seconds, and extension at 72^0^C for 2 minutes. After a final extension step at 72^0^C for 5 minutes, samples were run on a 2% agarose gel. The gels were imaged on Bio-Rad ChemiDoc, and the quantification for the band intensities was performed with the ImageLab software. The splicing efficiency of the respective genes was calculated by dividing the band intensities of the inclusion by those of the exclusion product. The values were further normalised with the respective eGFP condition as well.

### Microscopy analysis

After the growth in respective media, yeast cells were centrifuged at 14000 rpm for 15 seconds, and pellets were resuspended in 10µl of media. A total of 5µl of the cell suspension was spotted on a coverslip for live cell imaging. The Deltavision Elite microscope system was used to acquire all the images. The system was equipped with softWoRx 3.5.1 software (Applied Precision, LLC) and an Olympus 100x, oil-immersion 1.4 NA objective. The channel’s exposure time and transmittance settings were selected depending on protein expression and kept the same for all the biological replicates within an experiment. Images were captured as 512 × 512-pixel files with a CoolSnapHQ camera (Photometrics) using 1 × 1 binning for yeast. All the images were deconvolved using standard softWoRx deconvolution algorithms. ImageJ was used to analyze the data, and the granules were counted using the ‘Find Maxima’ tool from Fiji-ImageJ software. The images were converted to 8-bit, and the plugin was run. The prominence was set from 10-30, and the number of condensates and cells was counted.

The microscopy image acquisition for mammalian cells was performed using the Andor Dragonfly Confocal Microscope or Leica SP8 Falcon Confocal Microscope (for live cell peptide uptake experiment). HEK293T cells were imaged using a 63X objective, and the exposure time and transmittance were adjusted according to the protein expression levels and were kept the same for all the biological replicates within an experiment. Analysis was carried out using Fiji-ImageJ software.

Total fluorescent intensities or fraction of protein in condensates were calculated by measuring the CTCF (Corrected Total Cell Fluorescence) values for the ROI. For background subtraction, three regions from the background were selected, and the intensity was calculated from all three regions. This was followed by subtraction of background from the total intensity of ROI (region of interest) using the following formula:

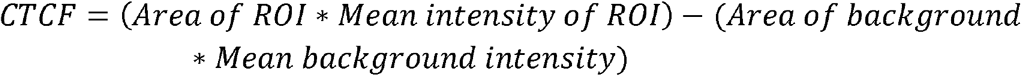

For fraction condensate intensity, the intensity of all condensates from a cell is quantified and divided by that of the total cell intensity. For cells with no condensates in the recovery phase, the value was kept at 0.

For nuclear: cytoplasm ratio analysis, the CTCF of the nucleus and the total cell were quantified by the aforementioned method. The cytoplasm intensity was calculated by subtracting the nuclear intensity from that of the total cell intensity. This was followed by calculating the ratio of N: C intensity. To normalize the values (for cells without condensates and N: C ratio in Figure 2) for the mScarlet expression levels, the respective values were divided by the mean CTCF mScarlet values.

### Expression and purification of recombinant proteins

All the primary cultures of *E coli* expression strains were cultured in LB and were grown at 37^0^C overnight in the presence of appropriate antibiotics. The individual protocols for the recombinant expression and purification of the proteins used in this study are detailed below. After purification, the elutes were dialyzed against the respective buffers, flash frozen, and stored at -80^0^C as 50-100µl aliquots.

### Purification of MBP-TeV-FUS-eGFP-TeV-His and MBP-TeV-FUS-Flag-TeV-His

MBP-TeV-FUS-eGFP-His and MBP-TeV-FUS-Flag-TeV-His expressing pMal plasmids were transformed into *E. coli* BL21 (DE3) Rosetta competent cells, which were selected on 100µg/µl ampicillin and 50 µg/µl chloramphenicol containing LB agar plates. A single colony was inoculated in LB media (containing 100µg/µl ampicillin and 50 µg/µl chloramphenicol) and was incubated overnight at 37^0^C and 180rpm. A secondary culture was set up from the primary culture and was grown to a final OD_600_ of 0.8. The cultures were then subjected to a cold shock by incubating the flasks on ice for 15 minutes. This helps in the induction of chaperones that help in preventing the aggregation of FUS. Post-cold shock, the FUS protein was induced with 1mM IPTG and was incubated at 10^0^C for 24 hours. Following induction, the cells were pelleted at 4200rpm at 4^0^C for 15 minutes. The cell pellets were stored at - 80^0^C.

For the Ni-NTA purification of the His tagged FUS, the cells were first resuspended in the resuspension buffer containing 50mM NaH_2_PO_4_, pH 8.0, 300mM NaCl, 10mM ZnCl_2_, 40mM imidazole,4mM beta-mercaptoethanol and 10% glycerol by vortexing. To the completely resuspended cells, a final concentration of 1mg/ml RNAase,1mg/ml lysozyme, and 1X PIC was added, followed by 30 minutes of incubation. Cells were lysed by sonication at 40% amplitude for 10 minutes with 10s on and off cycles. The lysate was centrifuged at 15000rpm for 15 minutes, and the supernatant was collected in a fresh tube. Ni-NTA resin calibrated with the resuspension buffer was added to the supernatant and was incubated for binding for 2 hours at 4^0^C in a nutator. After binding, the beads were washed thrice with the wash buffer (resuspension buffer without glycerol) and eluted with 500mM imidazole. The beads were spun down at 1500rpm for 1 minute, and the elute was collected. The FUS elute (this double step was used only for MBP-TeV-FUS-eGFP-His) obtained from the Ni-NTA purification was then subjected to binding to MBP resin, which was calibrated with the resuspension buffer overnight at 4^0^C in a nutator. After binding, the MBP beads were washed twice with the resuspension buffer and were eluted using resuspension buffer with 20mM maltose. The eluted proteins were then dialyzed using a buffer containing 20mM NaH_2_PO_4_, pH 8.1, 150mM NaCl, 5% glycerol, 1mM EDTA, and 1mM DTT. The dialyzed protein concentrations were measured using Bradford, followed by flash freezing and storage at - 80^0^C^43^.

### Purification of TeV protease

TeV protease was expressed in *E coli* BL21 (DE3) Rosetta pLys competent cells grown in standard LB media. TeV protease was induced overnight (16h) with 1mM IPTG at 0.6 OD_600_ at 20^0^C. The induced cell pellets were resuspended in Tris lysis buffer (50mM Tris pH8, 200mM NaCl, 20mM imidazole, 10% glycerol, and 4mM b-mercaptoethanol) and were supplemented with 1mg/ml RNAase,1mg/ml lysozyme, and 1X PIC. The resuspended cells were lysed by sonication. The TeV-His in the supernatant was purified by using Ni-NTA beads and washed with the Tris lysis buffer with 1M NaCl. The TeV protease was eluted using 800mM imidazole and was dialyzed against storage buffer containing 20mM Tris pH 7.4, 150mM NaCl, 20% glycerol, and 2mM DTT^43^.

### Purification of His-Sbp1-Flag, His-Sbp1ΔRGG-FLAG, His-Sbp1-AMD-Flag, His-Sbp1-GFP, and His-Sbp1ΔRGG-GFP

Recombinant His-Sbp1-Flag, His-Sbp1ΔRGG-FLAG, His-Sbp1-AMD-Flag, His-Sbp1-GFP, and His-Sbp1ΔRGG-GFP transformed in *E. coli* BL21 cells were induced with 1mM IPTG for 3 hours at 37^0^C. The induced cell pellet was resuspended in lysis buffer (300mM NaCl, 50mM NaH_2_PO_4_, 1mM DTT, 1mg/ml RNase, 1mg/ml lysozyme, 1X PIC, and 10mM Imidazole) followed by sonication. Lysate was clarified by centrifugation at 15000rpm for 15 minutes at 4^0^C. Clarified lysate was incubated with equilibrated Ni-NTA resin for 2 hours at 4^0^C. The His-Sbp1-GFP beads were washed thrice with wash buffer (300mM NaCl and 50mM NaH_2_PO_4_) containing increasing concentrations of imidazole at each step (20, 35, and 50mM). Protein was then eluted in elution buffer (50mM NaH_2_PO_4_, 300 mM NaCl, and 500 mM Imidazole). The protein was dialyzed into dialysis buffer (10mM Tris pH 7.0, 100 mM NaCl, 10% glycerol, and 1mM dithiothreitol) overnight at 4^0^C. The concentration was checked by Bradford assay, and small protein aliquots were stored at -80^0^C until further usage.

### In-cell sedimentation assay

HEK293T cells (from one T75 flask) transfected with the respective plasmids were collected 24 hours post-transfection. The cell lysis was carried out in RIPA buffer (50mM Tris-HCl, 1% NP-40, 150mM NaCl, 1mM EDTA, 1mM Na-orthovanadate, 1mM Na-fluoride, 1X PIC, 1mM PMSF, and RNase Inhibitor) for 30 minutes at 4^0^C. Cell debris was removed by a mini-spin at 1000g for 2 minutes at 4^0^C. Protein concentration was estimated using Bradford, and 700-800ug of lysate was further taken to separate the soluble and insoluble phases from the lysate by centrifugation at 15000g for 15 minutes at 4^0^C. The supernatant was collected as the soluble cytoplasmic fraction in a fresh tube, and the pellet was resuspended in 100ul of RIPA buffer. The disassembly reaction was set up in a fresh tube with 20ul of the pellet fraction and 5uM of the purified protein. This was allowed to incubate at 30^0^C for 1 hour and then centrifuged at 15000g for 15 minutes at 4^0^C to separate the soluble and the pellet fractions. The pellet was resuspended in 2% SDS buffer (2% SDS, 100mM Tris-HCl, pH 7.0) and incubated for 1 hour at 30^0^C. The resulting solution after that was considered to be the insoluble fraction. All samples were heated at 100^0^C for 5 minutes in 1X SDS loading dye and were taken ahead for Western analysis.

### *In vitro* sedimentation assay for FUS and TDP43

For the *in vitro* sedimentation assay of FUS, the phase separation of the purified MBP-FUS-eGFP-His was induced by the cleavage of the MBP tag by the TeV protease. A 50µl phase separation mixture with 1µM FUS in a buffer containing 50mM Tris pH 8, 0.5mM EDTA, and 1mM DTT was incubated at 30^0^C for 1 hour with the addition of 0.1mg/ml TeV protease^43^. A change in turbidity after 1 hour indicates phase separation. The test proteins were added at the mentioned concentration to FUS in the phase separation reaction to assess their impact on phase separation. After 1 hour of phase separation, the samples were centrifuged at 20000g for 15 minutes, and the soluble and insoluble fractions were separated. The insoluble (pellet) fraction was resuspended in 50µl buffer, and equal volumes of the two fractions were loaded onto a SDS-PAGE gel. CBB staining of the SDS-PAGE gels was carried out, and the ratio of FUS in the soluble to the total protein loaded (sum of FUS in the soluble and insoluble) was calculated from the band intensities analyzed by ImageLab software. The same *in vitro* sedimentation assay protocol was also used to assess the effect of Sbp1 on TDP43, with BSA as the negative control. The only difference in the case of TDP43 was the composition of the phase separation buffer. The MBP tag cleavage of TDP43 was carried out in a 50µl reaction mix with 1µM TDP43, HEPES buffer containing 20mM HEPES, pH 7.5, 150mM NaCl, and 1mM DTT with the addition of 20µg/µl of TeV.

### Interaction between FUS and Sbp1

For RFP-trap pulldown, HEK293T cells were co-transfected with FUS-P525L and Sbp1-related constructs. The cells were collected 48 hours after transfection, and cell lysis was carried out in RIPA buffer. Cell debris was removed by a mini-spin at 2000g for 2 minutes at 4^0^C. Protein concentration was estimated using Bradford, and 800-1000ug of lysate was further taken for the experiment. The lysate was diluted 1:1 with the dilution buffer (10mM Tris pH 7.5, 0.5mM EDTA, 150mM NaCl, 1X PIC, 1mM PMSF, and 0.05U/ul of Pierce Universal Nuclease), and 10µl of RFP-TRAP magnetic agarose beads (Cat. No. rtmak) were added to a final volume of 500ul. The pull-down samples were nutated at 4^0^C for 120⍰minutes. The beads were then separated using a magnetic stand, followed by two washes with the dilution buffer. Finally, the beads were resuspended in 40µl of dilution buffer, and Western analysis was carried out.

For *in vitro* interaction, purified MBP-FUS-Flag-His (0.1uM), His-Sbp1-GFP (0.1uM), and His-Sbp1ΔRGG-GFP (0.2uM) was used. An increased amount of His-Sbp1ΔRGG-GFP was used to compensate for the lower detection sensitivity of the anti-GFP antibody for this protein, ensuring levels comparable to His-Sbp1-GFP. The purified MBP-FUS-Flag was added to a pulldown reaction with either buffer or Sbp1-GFP/Sbp1ΔRGG-GFP. No bait control was set up for both Sbp1-GFP and Sbp1ΔRGG-GFP without FUS-Flag. The pull-down reaction contained purified protein, 1X dilution buffer (50 mM Tris pH 7.5, 150mM NaCl, 1mM EDTA, 1% Triton X-100) and 0.08% BSA to a final volume of 500 µl. 40ul of input sample was collected at this stage and 10ul of FLAG agarose beads (pre-blocked overnight in 2% BSA) were added to each reaction. The pull-down samples were nutated at 4^0^C for 2 hours. Subsequently, the beads were first washed with dilution buffer containing 150mM NaCl followed by 2 more washes with dilution buffer containing 500mM NaCl at room temperature for 5 minutes each. Finally, the beads were resuspended in 40ul of dilution buffer. Western analysis was performed to detect pulldown of target and bait protein.

### FUS self-association

To assess the self-association, cell lysates were prepared from HEK293T cells transfected with the eGFP-FUS-P525L expressing vector as described above. The reaction was set up and included 800-1000ug of lysate, 0.2uM of recombinant MBP-FUS-eGFP, and 300ul of the dilution buffer (supplemented with 1X PIC, 1mM PMSF, and 0.05U/ul of Pierce Universal Nuclease). The reaction was incubated at 4^0^C for 601minutes. This was followed by the addition of 0.2uM recombinant Sbp1 or Sbp1ΔRGG to the respective tubes. 20ul of input sample was collected at this stage, and the rest of the reaction was further incubated for 1 hour in a nutator at 4^0^C. MBP-sepharose beads were then added and allowed to nutate again for another 4^0^C. Finally, the beads were washed with dilution buffer three times and ultimately resuspended in 40ul of dilution buffer. Western analysis was carried out to check the immunoprecipitation of target and bait proteins.

For *in vitro* competition assay, 0.1uM of 2 differently tagged FUS variants (MBP-FUS-Flag-His and MBP-FUS-eGFP-His) were used. First, the MBP tag was cleaved from MBP-FUS-Flag-His using TeV protease in 1X Flag pulldown dilution buffer (50mM Tris pH 7.5, 150mM NaCl, 1mM EDTA, 1% Triton X-100) at 30^0^ C for 45 minutes. The cleaved FUS-Flag was then added to the FLAG pulldown reaction mixture containing 1X dilution buffer, 0.08% BSA and 10ul of FLAG agarose beads (pre-blocked overnight in 2% BSA) to a final volume of 500ul. The pull-down samples were nutated at 4^0^C for 60 minutes. This was followed by the addition of cleaved FUS-eGFP and either 0.1uM Sbp1-GFP or dialysis buffer to the respective pulldown reaction. 20ul of input sample was collected at this stage, and the rest of the reaction was further incubated for 60 minutes in a nutator at 4^0^C. Subsequently, the beads were first washed with dilution buffer containing 150 mM NaCl followed by 2 more washes with dilution buffer containing 500mM NaCl at room temperature for 5 minutes each. Finally, the beads were resuspended in 40ul of dilution buffer. Western analysis was performed to detect pulldown of target and bait protein. The amount of FUS-eGFP normalized to FUS-Flag pulldown was used as a quantitative measure of FUS self-association.

### Western analysis

Western analysis was conducted as per the standard protocol. Blocking was done using 5% skim milk powder in TBST. Primary antibody incubation was done overnight at 4^0^C. The blots were incubated at room temperature for 1 hour with the secondary antibody and were developed using the Western ECL kit in a Bio-Rad ChemiDoc. The antibodies used in this study are anti-GFP (Biolegend, 902602), anti-Pgk1 (Abcam, Ab113687), anti-Gapdh (Cloud-clone corp., MAB932Hu23), anti-mCh (Abcam, Ab167453), anti-flag (Sigma, F3165), anti-rabbit (Jackson ImmunoResearch Lab, 111-035-003), anti-mouse (Jackson ImmunoResearch Lab, 115-035-003), and anti-rabbit-alexafluor568 (Invitrogen, A11011).

### Statistical analysis

All the analyses were conducted using the GraphPad Prism software (version 8.0.2). The significance was calculated either by unpaired/paired Student t-test or Tukey’s multiple comparisons test (2-way ANOVA). All the figure legends include the specific details of the test used and the p-value considerations. In all the graphs, the error bars represent the standard error of the mean (SEM), and the same colour points in a graph depict the data from a single experimental set.

## Acknowledgment

All the members of the Rajyaguru lab are acknowledged for their constant inputs, support, and encouragement. We thank the Indian Council of Medical Research (ICMR, grant#IIRPIG-2024-01-00233), Amyotrophic Lateral Sclerosis (ALS) Association (Grant#26-SGP-761) and the Department of Biotechnology, Government of India (DBT, grant#BT/PR51975/BMS/85/23/2024) for supporting this research; the Department of Science and Technology (DST-FIST) India, and the Indian Institute of Science (IISc) for infrastructure and other support. KS acknowledges the Indian Council of Medical Research (ICMR, grant#2021-14148/CMB/ADHOC-BMS) and Department of Biotechnology (DBT, grant#BT/PR50450/MED/12/1044/2023). MG thanks CSIR-UGC and SERB-India, NV thanks PMRF, and PG appreciates GATE for the financial assistance. Flowchart or model figures were generated from adapted images provided either by Biorender or Servier Medical Art (Servier; https://smart.servier.com; licensed under a Creative Commons Attribution 4.0 Unported License).

## Author contributions

Conceptualization and hypothesis—PIR and MG; Experimental design—PIR, MG, NV, RD, KS, and PG; Experimentation—MG, NV, KN, RD, and PG; Data interpretation—PIR, MG, NV, KN, RD, PG, and KS; Manuscript writing (first draft)—MG, Subsequent draft review and editing— PIR and MG.

## Supplementary Figures

**Supplementary Figure S1:**
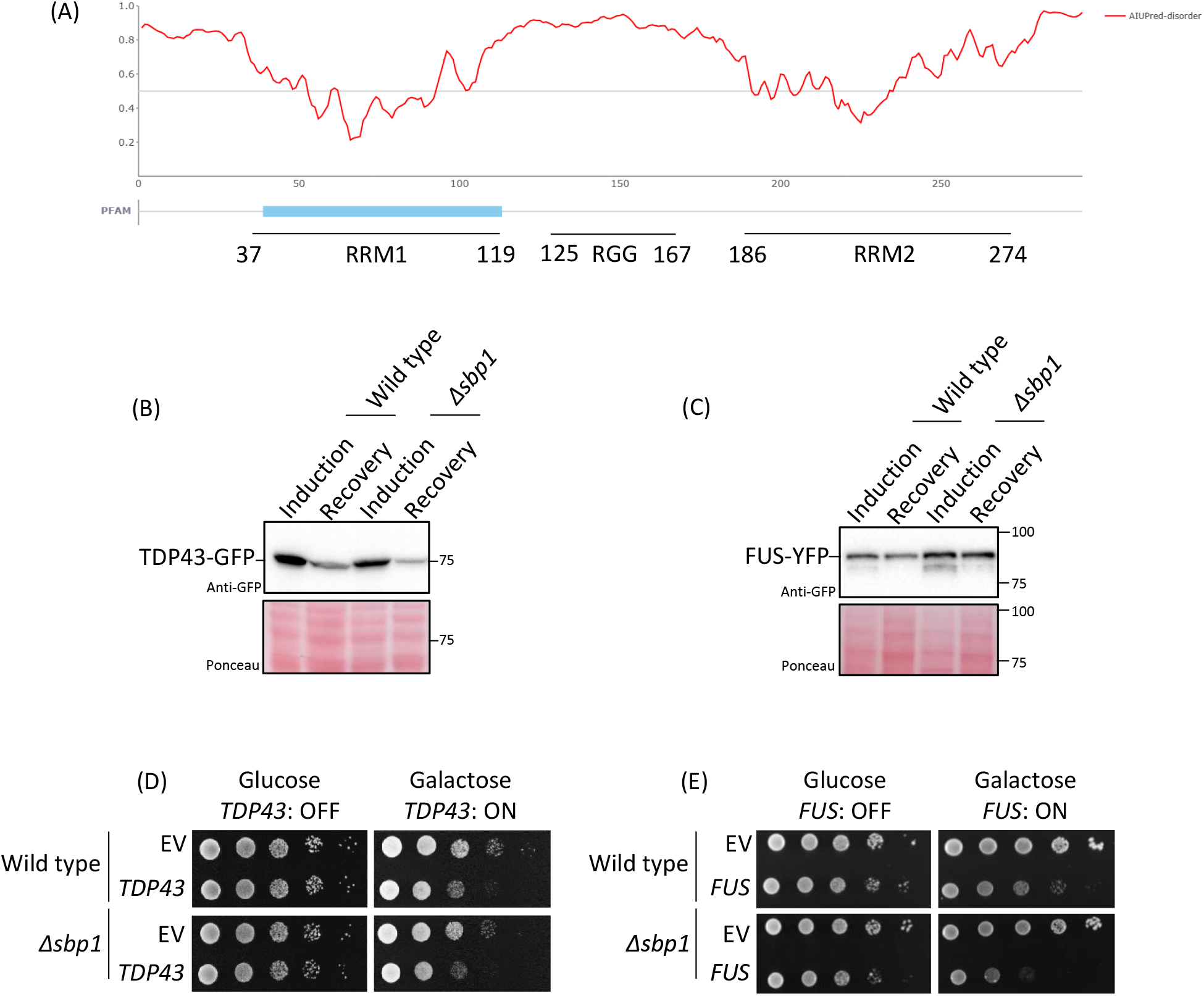
Δsbp1 cells are defective in the disassembly of TDP43 and FUS condensates in yeast. A) AIUPred-disorder analysis for Sbp1. Different domains/motifs are also marked. (B and C) Western analysis depicting the change in protein levels of TDP43 (B) and FUS (C) in the recovery state with respect to the respective induction condition. Quantitation of the blots is provided as Figure 1D and G. Values on the right represent the position of different molecular weight ladder bands in kDa. Ponceau served as the loading control. (D and E) Spot assays of wild type and Δ*sbp1* cells transformed with either empty vector (EV) or Gal-*TDP43-GFP* (D) / Gal-*FUS-YFP* (E) expressing constructs.

**Supplementary Figure S2:**
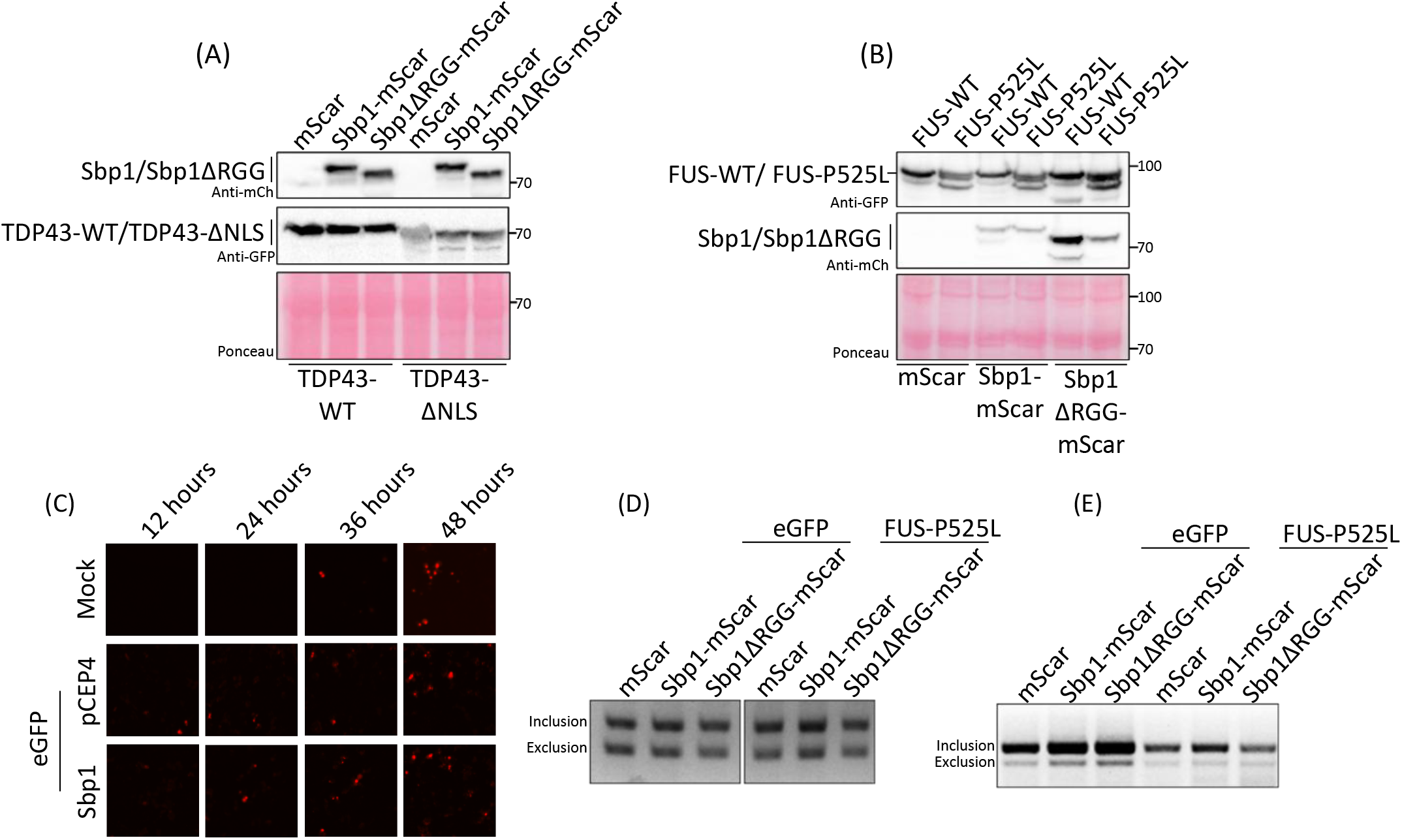
RGG-motif protein expression reduces mutant TDP43 and FUS condensates and overexpression-mediated defects of FUS-P525L in mammalian cells. (A and B) Western analysis to check the levels of TDP43-WT/ΔNLS (A) or FUS-WT/P525L (B) upon expression of Sbp1/Sbp1ΔRGG. Ponceau serves as the loading control, and the values on the right denote the position of different molecular weight ladder bands in kDa. Quantitation of the blots is provided as Figure 2C and F. (C) Incucyte images representing the cellular uptake of propidium iodide (PI) in different conditions in HeLa cells. Scale bar=100um. The images are part of Figures 2H and I. (D and E) Agarose gel images depicting the bands for the inclusion and exclusion products for *hnRNP-D* (D) and *TTC3* (E) RNAs from N2A cells expressing eGFP/FUS-P525L, along with either mScar/Sbp1/Sbp1ΔRGG. The ratio of the inclusion to exclusion was calculated and plotted in Figure 2L and M.

**Supplementary Figure S3:**
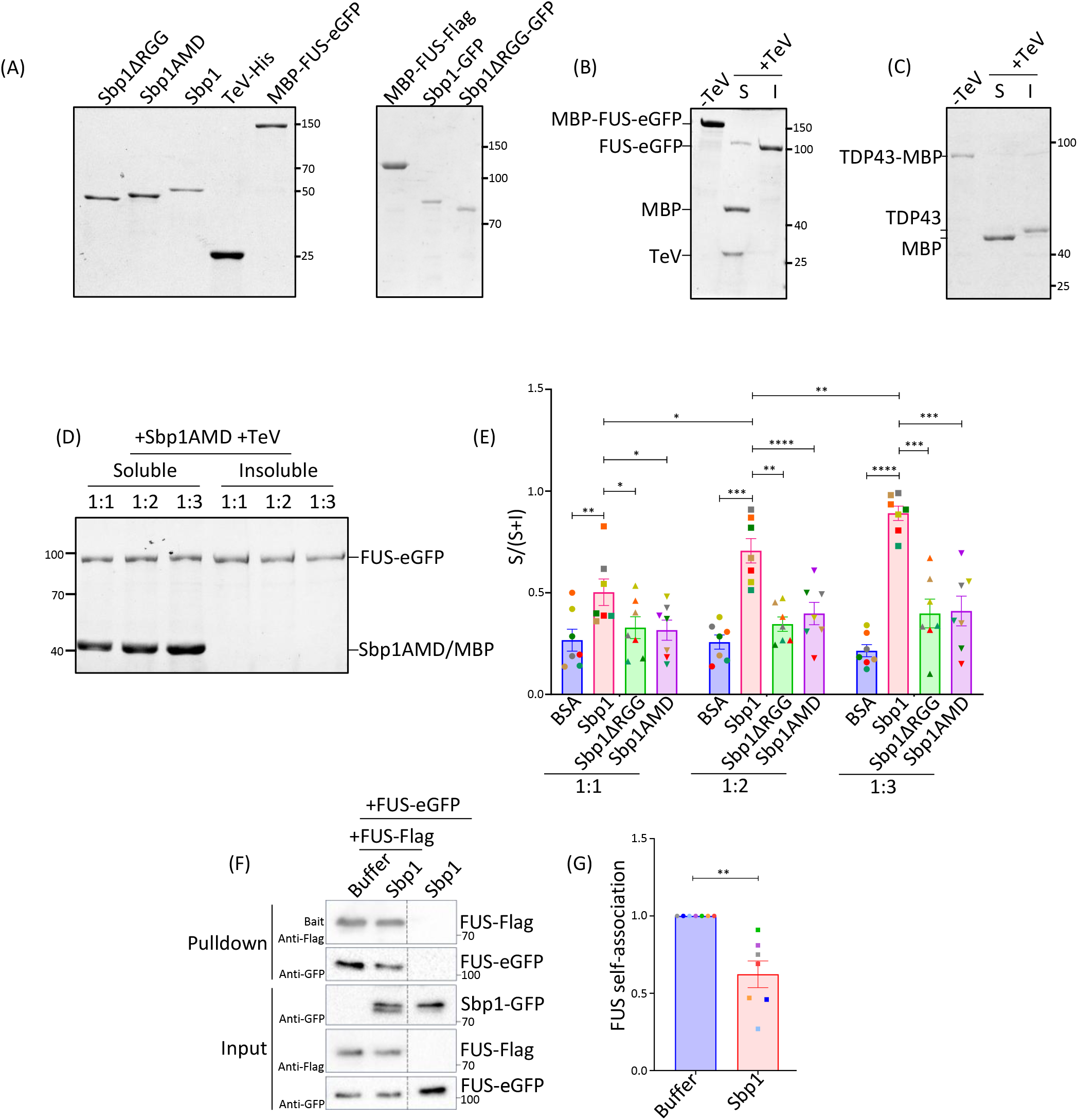
Sbp1AMD (arginine methylation defective) mutant fails to disassemble FUS condensates. (A) Coomassie-stained protein gel depicting the various purified proteins used in this study. Values on the right represent the position of different molecular weight ladder bands in kDa. (B and C) Coomassie-stained protein gels depicting the partitioning of FUS (B) and TDP43 (C) to the insoluble (I) phase after the cleavage of the MBP-tag using TeV protease. S denotes the soluble phase. Values on the right represent the position of different molecular weight ladder bands in kDa. (D) Coomassie-stained protein gel depicting the fractionation of FUS to soluble and insoluble phases in the presence of Sbp1AMD (mutant where all arginines within the RGG-motif are covered to alanine, see Figure 1A). The ratio reflects the amount of FUS: test protein taken for the assay. Sbp1AMD and MBP migrate at the same position and hence appear as a single band. Values on the left represent the position of different molecular weight ladder bands in kDa. (E) Quantitation of the fraction of FUS protein present in the soluble phase from 7 independent experiments (n=7) as performed in D and Figure 3J-L. The graph from Figure 3M has been replotted here to include Sbp1AMD values. Significance was calculated by Tukey’s multiple comparisons test (2-way ANOVA). (F) Western analysis depicting the interaction of FUS-eGFP upon pulldown of FUS-Flag in the presence or absence of Sbp1. Sbp1+FUS-eGFP lane had everything except the bait, FUS-Flag. Values on the right represent the position of different molecular weight ladder bands in kDa. Dashed line denotes some middle lanes that were cropped because these were not relevant for this figure. (G) Quantitation for the amount of FUS-eGFP from the pulldown experiment as performed in F. The values were normalised with respect to the amount of bait in the pulldown reaction. A student paired t-test analysis was used to calculate the significance (n=7). Error bars represent mean ±SEM, and the same color points depict the data from a single experimental set. *, **, ***, and **** denote p-value ≤0.05, ≤0.01, ≤0.001, and ≤0.0001, respectively.

**Supplementary Figure S4:**
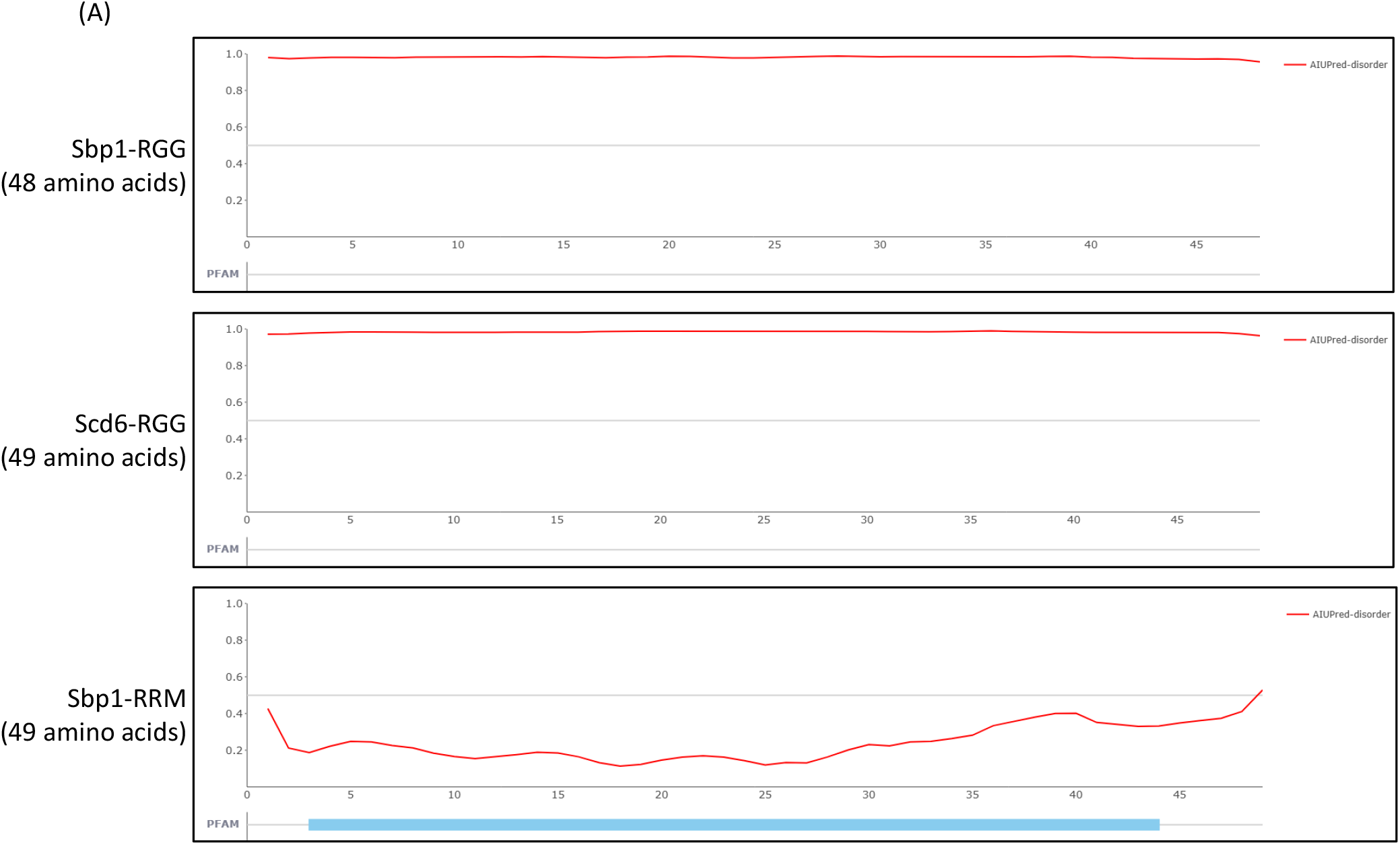
Disorder analysis of the different peptides. (A) AIUPred-disorder analysis for the different peptides used in Figure 4.

